# Assembly status transition offers an avenue for allosteric activity modulation of a supramolecular enzyme

**DOI:** 10.1101/2021.08.20.457084

**Authors:** Yao Chen, Weiya Xu, Shuwei Yu, Kang Ni, Guangbiao She, Xiaodong Ye, Qiong Xing, Jian Zhao, Chengdong Huang

**Author notes:** These authors contributed equally (Yao Chen, Weiya Xu, Shuwei Yu). Corresponding author: Qiong Xing; Jian Zhao; Chengdong Huang.

## Abstract

Nature has evolved many supramolecular proteins assembled in certain, sometimes even seemingly oversophisticated, morphological manners. The rationale behind such evolutionary efforts is often poorly understood. Here we provide atomic-resolution insights into how the dynamic building of a structurally complex enzyme with higher-order symmetry offers amenability to intricate allosteric regulation. We have established the functional coupling between enzymatic activity and protein morphological states of glutamine synthetase (GS), an old multi-subunit enzyme essential for cellular nitrogen metabolism. Cryo-EM structure determination of GS in both the catalytically active and inactive assembly states allows us to reveal an unanticipated self-assembly-induced dynamics-driven allosteric paradigm, in which the remote interactions between two subcomplex entities significantly rigidify the otherwise structurally fluctuating active sites, thereby regulating activity. We further show *in vivo* evidences that how the enzyme morphology transitions could be modulated by cellular factors on demand. Collectively, our data present an example of how assembly status transition offers an avenue for allosteric modulation, and sharpens our mechanistic understanding of allostery, dynamics, cooperativity, and other complex functional and regulatory properties of supramolecular enzymes.

## Introduction

Recent studies have evidenced that only a small portion of proteins function in isolation in cells whereas the majority is assembled into complexes through protein-protein interactions with identical or different protein subunit(s) ^1^. The rationale behind such an evolutionary selection has been the subject of considerable speculation; proposals for the advantages associated with a multimeric-units complex instead of a long single polypeptide chain include better error control in synthesis, greater coding and folding efficiency, and possibility of allosteric regulation ^2^. Morphologically speaking, many protein complexes especially homomeric ones adopt a symmetric spatial arrangement, either cyclic (C_n (n>1)_) or dihedral (D_n (n>1)_) symmetry, characterized by a rotational symmetry or two orthogonal symmetry axes, respectively. In contrast to the cyclic complexes which evolve in one step (e.g. C1→C5), evolution of dihedral complexes takes place in multiple steps (e.g. C1→C5→D5) ^3^, and adds another layer of structural complexity. Intriguingly, pioneering studies have revealed many supramolecular enzymes organized in dihedral symmetry, with subcomplex entities in cyclic symmetry holding, at least outwardly, multiple integral active sites. Thus a fundamental question arises here is that why nature builds these protein complexes with a seemingly oversophisticated quaternary design, if the subcomplexes alone possess complete elements for action? In other words, is the extra assembly step, e.g. C5→D5, is a futile evolutionary effort for these supramolecular protein complexes?

One such an example is glutamine synthetases (GSs) (EC 6.3.1.2), one of the most ancient functioning enzymes in existence and a central enzyme in nitrogen metabolism of all living organisms, catalyzing the formation of glutamine by condensation of glutamate with ammonia using ATP as an energy source ^4,5^. Three classes of GS enzymes have been identified in different organisms, namely, GSI, GSII and GSIII. Decades of studies have established a striking notion that all three classes of GS enzymes, despite of dramatic differences in amino acid sequences and protein sizes, share quaternary geometry in dihedral symmetry assembled with two oligomeric rings ^6–13^. Considering that the active sites of GS are located at the clefts formed between two neighboring protomers within the same ring and distal to the ring-ring interface, each isolated GS subcomplex ring holds multiple integral catalytic sites ^4,14^. The functional demand for this evolutionary conservation, the quaternary organization of GS with dihedral symmetry, remains elusive.

Here we sought out to explore the functional link between the oligomeric conformation and catalysis activity, and mechanistically justify the seemingly oversophisticated assembly design in this supramolecular enzyme. Our results unveil a previously uncharacterized dynamics-driven allostery mechanism induced by assembly status transition of GS, and present an example that how a particular quaternary geometry selectively defines the oligomer dynamics congruent with required allosteric activities. We further show *in vivo* evidence how this regulatory machinery is elegantly utilized by the cell to meet the ever-changing metabolic needs. The functional implications of these findings are discussed.

## Results

### Two highly conserved GSIIs demonstrate distinct quaternary structure organization propensities

With the aim of clarifying the functional role of dihedral symmetry in GS functions, we first carried out a quest for GSs that share a high degree of sequence conservation, but demonstrate distinct quaternary structural assembly properties. We make use of the weak ring-ring interaction of GSII, a prominent structural difference between the type I and type II GSs ^11–13^, and seek GSII variants from different species with amino acid variations mainly occurring at the pentamer interface, which may thus present disparate decamer-forming propensities. We built model structures for the candidate GSIIs based on the crystal structure of the maize GSII (pdb code:2D3B) and analyzed the amino acid variations in the context of model structures. Primary structure analysis reveals GSIIs from the plants of *Camellia sinensis* (CsGSIb) and *Glycine max* (GmGSβ2) share an overall very high sequence homology (∼90% identical and ∼97% conserved) and absolutely conserved substrate-binding and catalytic sites (Fig. 1a), with, however, a significant portion of amino acid variations clustered at the interface between two pentamer rings (Fig. 1a and 1b). We then recombinantly expressed both CsGSIb and GmGSβ2 in *E. coli* and purified these two GSII homologs. To assess the oligomerization status, we performed size-exclusion chromatography (SEC) coupled to both multi-angle light scattering (MALS) and quasi-elastic light scattering (QELS). Despite sharing an overall highly conserved amino acid sequence, CsGSIb and GmGSβ2 exhibit distinct quaternary structural properties. MALS analysis shows the GSII from *Glycine max* being largely a homogeneous decamer in solution (Fig. 1c). In contrast, under the same condition the majority fraction (∼82%) of CsGSIb adopts a pentameric configuration, along with a minor fraction (∼18%) being decameric (Fig. 1d). We further show that CsGSIb exists in pentamer-decamer dynamic equilibrium in solution and a mixture of electrostatic and hydrophobic interactions is responsible for attaching of two pentameric rings; whereas substrates or ligands show no appreciable effect on the decamer-forming properties (see Supplementary data for details).

**Fig. 1:**
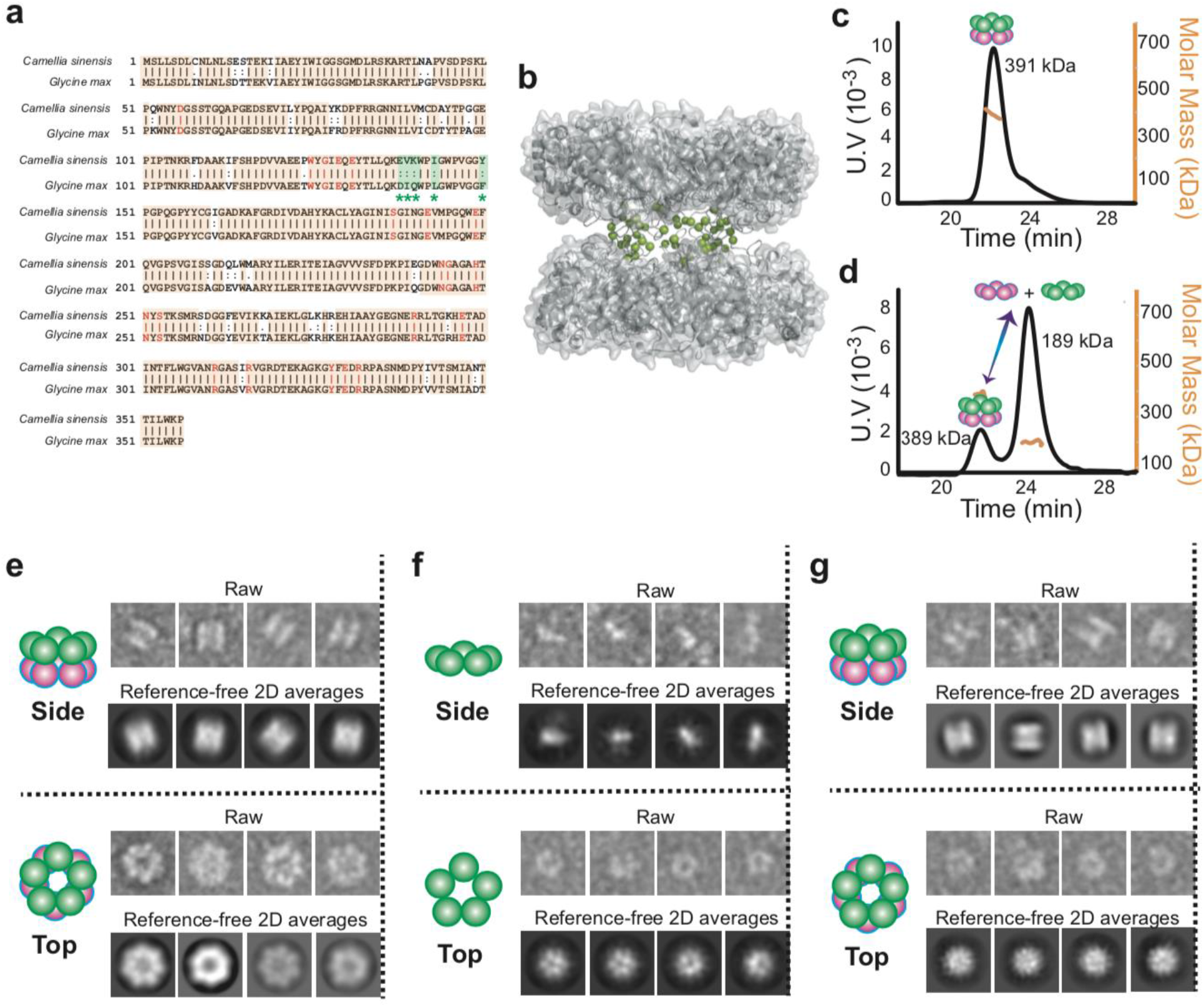
Quaternary assembly property comparison of GSIIs from *Camellia sinensis* (CsGSIb) and *Glycine max* (GmGSβ2). **a**, Amino acid sequence alignment of CsGSIb and GmGSβ2 reveals very high level of conservation. Identical amino acids are shown with orange boxes, while the residues involved in substrate-binding and catalysis are shown in red. Amino acids variations located at the pentamer ring-ring interface are highlighted in green boxes with the symbol of *. **b**, Model structure built based on the crystal structure of a maize GSII (GmGSβ2, pdb code 2D3B). The amino acid variations between CsGSIb and GmGSβ that are located at the ring-ring interface are highlighted as spheres in green. **c-d**, SEC-MALS analysis of GmGSβ2 (c) and CsGSIb (d). **e-g**, Quaternary assembly analysis of GmGSβ2 and CsGSIb using negative-stain electron microscopy. Left: Examples of single raw images; Middle: reference-free two-dimensional class averages. Right: A schematic representation of the averages is shown for clarity. GmGSβ2 adopts a homogenous double-ringed structure (e), while the CsGSIb demonstrates a mixture of two major classes of particles: isolated pentamer ring (f) and double-ringed structure (g).

We further employed negative-stain electron microscopy (EM) to directly visualize the distinct ring-ring packing propensities for CsGSIb and GmGSβ2. Two-dimensional (2D) class averages revealed that GmGSβ2 forms homogeneous, double stacked-ring shaped particles (Fig. 1e), in line with the decameric organization pattern previously reported for other GSII species ^11–13^. In contrast, CsGSIb adopted a mixture of two quaternary structural modes: detached pentamers (Fig. 1f) and decamers composed of two stacked pentamer rings (Fig. 1g), consistent with the above MALS analysis result (Fig. 1d).

Analytical ultracentrifugation (AUC) was carried out to quantitively assess the thermodynamic parameters of CsGSIb pentamer-decamer transition, which demonstrated two species for CsGSIb in solution with molecular weights of 191 kDa and 395 kDa (Fig. 2a), corresponding to the pentameric and decameric configurations, respectively. Sedimentation profiles at various protein concentrations were analyzed and a global analysis of the data at each protein concentration yielded a pentamer-pentamer dissociation constant (Kd) of 0.27±0.06 μM at room temperature. It is noteworthy that the dissociation constant within the sub-micromolar range allows CsGSIb to predominantly exist as isolated pentamers under the concentration assayed for enzymatic activities, laying a solid foundation for probing the functional role of the decamer formation in modulating the enzyme activity of GS.

**Fig. 2:**
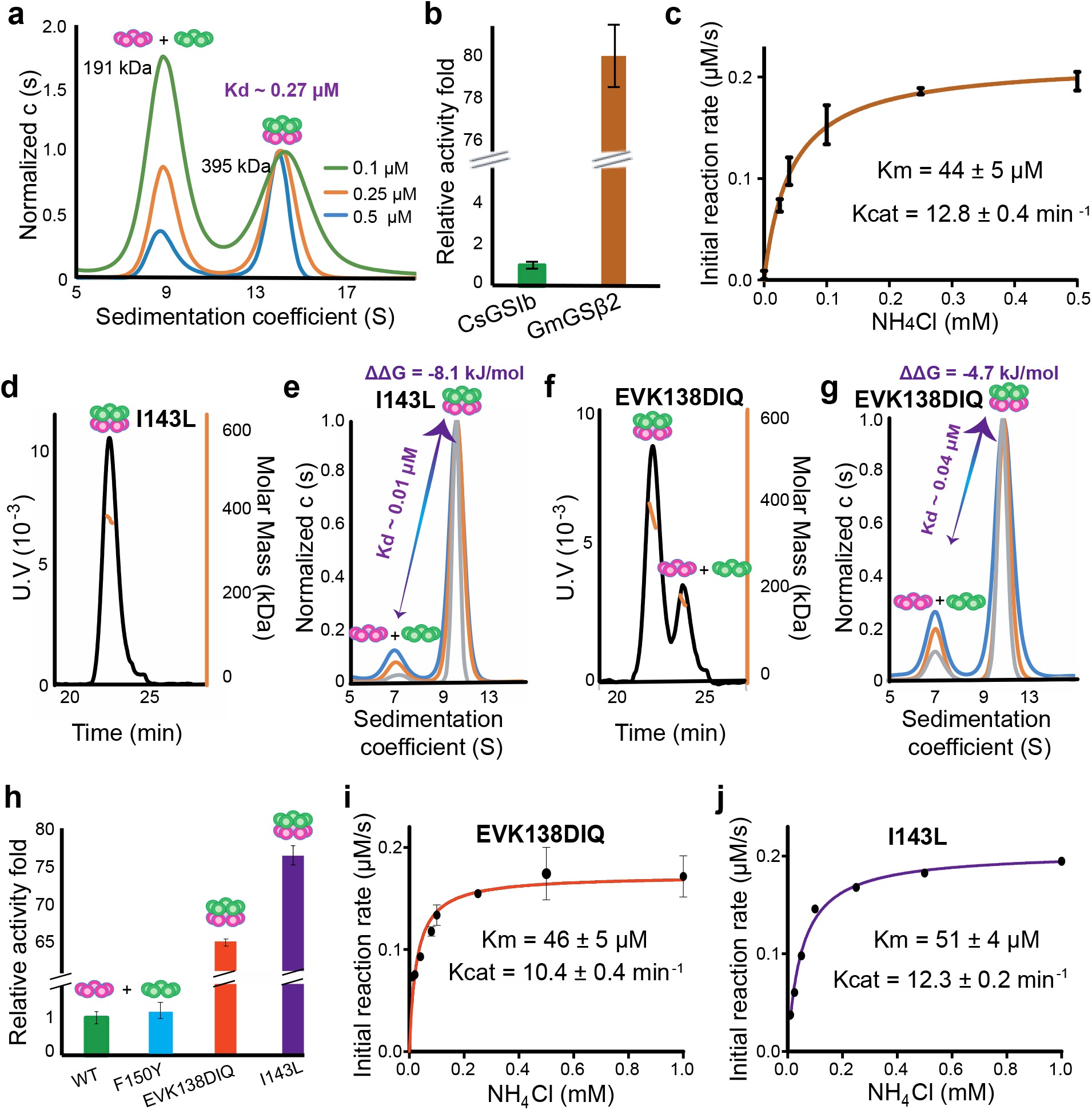
The enzymatic activities of CsGSIb is dependent on its quaternary assembly status. **a**, Application of analytical ultracentrifugation (AUC) to assess the pentamer-decamer dissociation constant of CsGSIb. Experiments were performed at room temperature with three different sample concentrations shown as monomer concentration. A global fit of sedimentation distribution profiles yielded a dissociation constant of 0.27 μM. **b**, Enzymatic activity comparison of GmGSβ2 with CsGSIb. Reactions were performed for 30 min at 37°C in presence of 1 μM (monomer) enzyme and saturated amounts of substrates. **c**, Steady-state kinetic analysis of GmGSβ2. Assay conditions were the same as that in **b**, except the concentrations of NH_4_Cl were varied. **d-g**, Mutation effects on the quaternary assembly property of CsGSIb evaluated using SEC-MALS (**d** and **f**) and AUC (**e** and **g**). The corresponding mutants are labeled in the figures. **h**, Activity comparison of the wild type GmGs18 with its mutants as labeled. Reactions were performed in the same condition as **b**. **i**-**j**, Steady-state kinetic analysis of GmGSβ2 mutants of EVK138DIQ (**i**) and I143L (**j**). Reaction conditions were same as in **c**. All enzyme assays were repeated at least three times and data were shown as means ± s.d.

### GSIIs in different assembly states demonstrate distinct enzymatic activities

We next sought to compare the glutamine synthesis activity of these two highly conserved GSIIs. As shown in Fig. 3b and Fig. S2a, when supplied with ammonium chloride, the stable decamer-adopting GmGSβ2 demonstrated significant GS activities. Further steady-state kinetic measurements yielded turnover numbers (k_cat_) and Michaelis constants (K_m_) of ∼12.8 min^−1^ and ∼44 μM, respectively (Fig. 3c). In sharp contrast, CsGSIb, for which the majority protein exists as discrete pentamers under the condition assayed, only demonstrated basal activity (Fig. 3b and Fig. S3a), consistent with previous observations that the isolated single-ringed GSII species is nonfunctional ^15,16^. These observations raised the question as to whether the drastic disparity in catalytic activities for these two highly conserved GSII could be attributed to their dramatically different propensities for formation of a double-ringed architecture. If the above proposal holds true, a positive concentration-dependent cooperation of enzyme activity would be expected as increase in the concentration of CsGSIb favors decamer assembly (Fig. S1a). Indeed, five times increase in the concentration of CsGSIb assayed (from 1 μM to 5 μM) resulted in ∼30 folds increase in activity, i.e. ∼6 folds activity increase per unit of enzyme, displaying a significant concentration-dependent simulation effect (Fig. S2b).

**Fig. 3:**
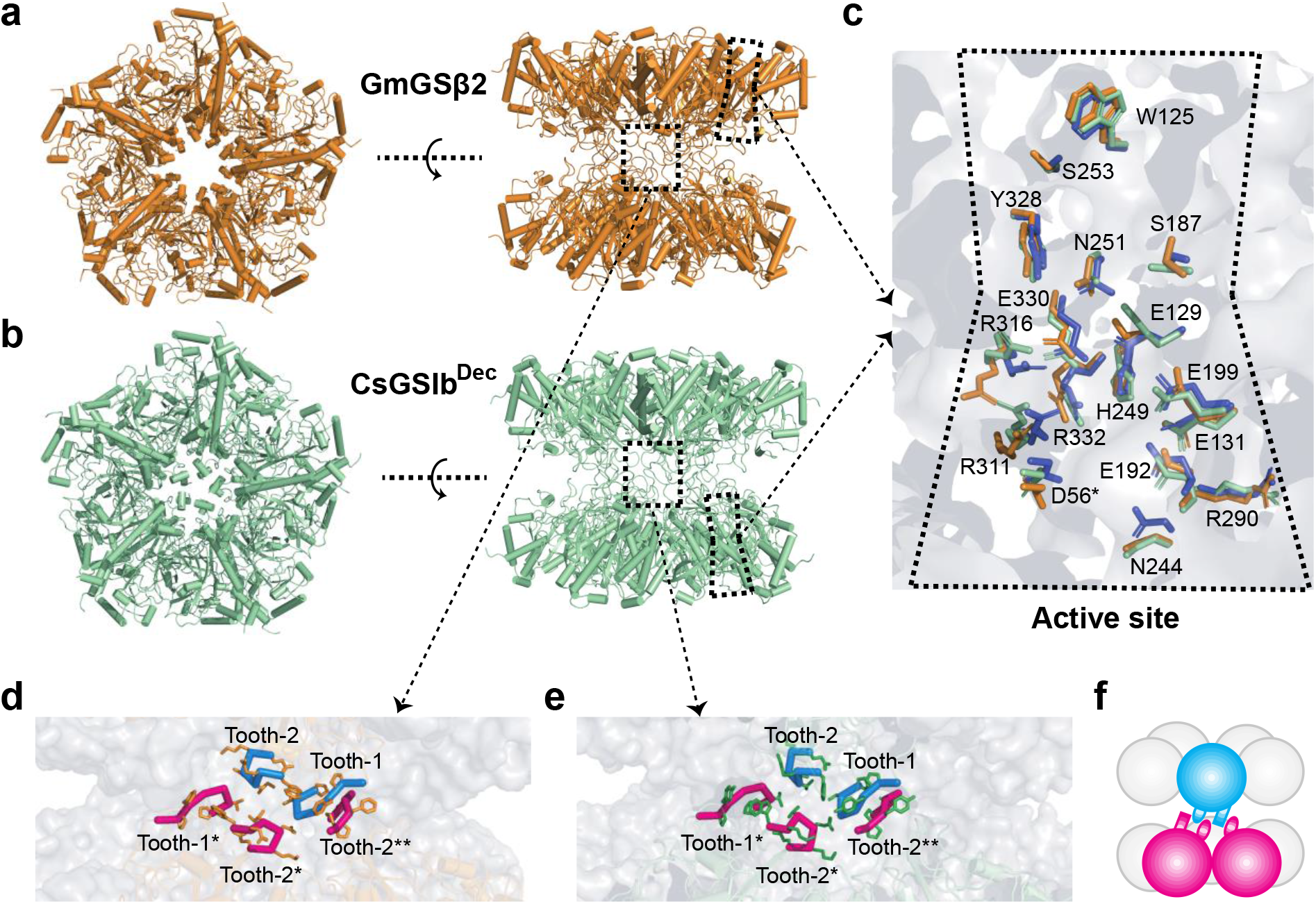
Overall structures, catalytic centers and ring-ring interfaces of GmGSβ2 and CsGSIb^Dec^. **a**-**b**, Overall double-ringed structures of GmGSβ (**a**) and CsGSIb^Dec^ (**b**). Left: top-view; Right: Side-view. **c**, Superimposed structures of the catalytic centers of GmGSβ2 (orange), CsGSIb^Dec^ (green) and GSII from maize (purple). High structure similarities in catalytic sites suggest the catalytic mechanism for these three GSII species are essentially identical. **d**-**e**, The detailed ring-ring interaction interfaces between GmGSβ2 (**d**) and CsGSIb^Dec^ (**e**). The interactions between two pentameric rings are primarily mediated by two regions, namely, the tooth-1 and tooth-2, respectively. (**f**), A schematic representation of the two GSII pentameric rings interlocked by tooth-1 and tooth-2 for clarity.

### Catalytic switching of CsGSIb through oligomeric states interconversion

To further validate the above proposal, we performed mutagenesis to CsGSIb, aiming to convert its unstable pentamer-decamer equilibrium state to a stable decamer and then evaluate the impact on catalytic activity. According to the amino acid sequence of GmGSβ2, three CsGSIb mutants, i.e. EVK138DIQ, I143L and Y150F, respectively, were selected, expressed and subsequently purified as the wild type (WT) protein. We then performed SEC-MALS measurements to evaluate mutational effects on their oligomeric states. As shown in Fig. 2dx and 2f, both mutations of I143L and EVK138DIQ led to a drastic shift in the oligomerization equilibrium towards decamer assembly, causing significant increase in the distribution of decameric state from ∼18% in the WT-CsGSIb (Fig. 1d) to >95% and ∼72%, respectively. These observations confirmed that both mutations introduced at the interface, albeit largely conservative, dramatically fortified the decamer edifice assembly. In contrast, substitution of the tyrosine at the residue 150 with a phenylalanine showed no appreciable change in its quaternary organization mode (Fig. S3a), suggesting no critical role for the residue Y150 in maintaining two CsGSIb pentameric ring subcomplexes attached. We employed AUC analysis to measure the ring-ring dissociation constant for mutants of I143L and EVK138DIQ. As demonstrated in Fig. 2e and 2f, replacement of I143 or EVK at residues 138-140 with leucine or DIQ yielded ring-ring disassociation constants of ∼0.01 or ∼0.04 μM, respectively, i.e., ∼27 or ∼7 folds of increase in the ring-ring binding affinity compared with that of WT-CsGSIb. The Gibbs energy changes upon mutation (ΔΔG) for I143L and EVK138DIQ were calculated to be -8.1 kJ/mol and -4.7 kJ/mol, respectively.

We next set out to investigate the resulting impacts on their enzymatic activities. As shown in Fig. 2h-2j, both mutations of I143L and EVK138DIQ caused dramatic increase in catalytic activity of ∼76 and ∼64 folds, respectively. As all mutated amino acids are distal to either the catalytic site or substrate binding regions with distances >20 Å and hence are unlikely to be directly involved in catalytic reaction, we infer that the stimulations of the enzymatic activity of CsGSIb upon residue perturbations are attributed to allosteric effects induced by remote contacts between two pentamer rings. As expected, the mutation of Y150F, which did not alter pentamer-decamer equilibrium of CsGSIb (Fig. S3a), showed no noticeable change in enzymatic activity (Fig. S3b).

### Structural basis for dynamics-driven allostery of GSII

To elucidate the allosteric mechanism of how the interactions between two GSII pentameric rings remotely trigger enzymatic activity, we next employed single-particle cryo-EM imaging technique and first determined the structures of GmGSβ2 decamer, as well as that of the CsGSIb that adopts decameric configuration (thereafter named as CsGSIb^Dec^). 3D classifications of 104717 and 43876 particles for GmGSβ2 and CsGSIb^Dec^, respectively, revealed that both molecules were arranged in D5 symmetry with two pentameric rings stacked in a head-to-head manner (Fig. 3a and 3b), a strikingly conserved structural feature that have been widely observed for other type II GS species ^11–13^. Refinement of the GmGSβ2 and CsGSIb^Dec^ structures yielded maps with an average resolution of 2.9 Å and 3.3 Å, respectively, with literally identical dimensions of 115 Å x 115 Å x 95 Å. As expected from the very high conservation in amino acid sequence (Fig. 1a), decamer structures of GmGSβ2 and CsGSIb^Dec^ are very similar to each other, as well as to that of the GSII of maize (pdb accession number 2D3A), as highlighted by the root-mean-square deviation (r.m.s.d.) of 0.66–0.80 Å for 328-352 aligned C_α_ atoms. Structural alignments reveal that the active sites in GmGSβ2 and CsGSIb^Dec^, as well as that in the maize GSII, are highly conserved (Fig. 3c), suggesting the catalysis mechanism of these three enzymes, once decamers are formed, are essentially identical. The overall buried inter-ring surfaces for both GmGSβ2 and CsGSIb^Dec^ amount to ∼2000 Å^2^, i.e. approximately only 400 Å^2^ per individual monomer–monomer interaction. This highlights the weakness of the inter-ring contacts, characteristic of type II GS. Indeed, in both structures the inter-ring contacts are established by only a limited number of hydrophobic and polar interactions provided by the residues 136-141 and 146-152 segments of each of the intervening subunits (Fig. S7), which behave as two gear teeth (thereafter named as tooth-1 and tooth-2, respectively) interlocking the two pentameric rings (Fig. 3d-f). This observation is in line with the above result that mutations to tooth-1 resulted in drastic change in oligomeric states behavior (Fig. 2d-2g), and the mixed nature of the inter-ring interactions is consistent with the MALS analysis result of CsGSIb under various buffer conditions (Fig. S1b). Intriguingly, the local structure of the teeth regions that mediate inter-ring interactions remains largely the same in GmGSβ2 and CsGSIb^Dec^ (Fig. 3d-3e), suggesting the dramatically different propensities of GmGSβ2 and CsGSIb for decamer formation are due to the nature of the amino acids involved in inter-ring contacts, rather than the structure. Although the residue of I143 is not directly involved in inter-ring contact, we argue that its replacement with leucine may stabilize the conformation of tooth-1 via its interaction with the residue of L134, thus playing an important role in stabilizing decamer architecture (Fig. 2d and 2e).

In order to elucidate the mechanism of how the pentameric CsGSIb (thereafter named as CsGSIb^Pen^) demonstrates distinct enzymatic properties than CsGSIb^Dec^ (Fig. 2h), we next determined the structure of CsGSIb^Pen^. Lowering the sample concentration, which favored pentamer dissociation (Fig. S1a), allowed us to obtain sufficient number of CsGSIb^Pen^ particles, which, in turn, enabled us to solve the cryo-EM structure of the inactive single-ringed GSII for the first time. Interestingly, we observed additional class averages in which the two masses of density attributed to the pentameric rings are no longer parallel (Fig. S8). These non•parallel ring particles may reflect intermediate assembly stages in the formation/disruption of the enzyme decamer, again confirming the flexibility of the inter-ring interactions of the type II GS.

Unexpected, CsGSIb^Pen^ exhibited high conformational heterogeneity and 3D particles classification generated three similar structures with the r.m.s.d. ranging from 0.6 to 0.8 Å, differing in a few peripheral regions (Fig. S9a). The most striking difference between CsGSIb^Pen^ and CsGSIb^Dec^ is that several regions are missing in the electron density map of all three classifications of CsGSIb^Pen^ particles, with only 229 to 255 out of 356 residues electron densities in presence. As a result, CsGSIb^Pen^ demonstrated a decagram-shaped density map with a few regions missing at the rim (Fig. 4a and Fig. S9a), in sharp contrast to a pentagon-shaped map yielded by the CsGSIb^Dec^ particles (Fig. 4b). For the 229-255 residues that show clear density in CsGSIb^Pen^ particles, the conformation of each subunit in three CsGSIb^Pen^ EM structures, as well as the arrangement pattern, closely resembles that of the CsGSIb^Dec^, as evidenced by r.m.s.d. in the range of 0.7-1.0 Å (Fig. S9b-d). This result suggest the structures of rigid portion of CsGSIb are not significantly altered upon pentamer association. The density-missing regions include the segments around residues of 110-117, 140-166, and 260-334, among which, the fragment around residues 260-334 is a major component making up an integral catalytic site (Fig. 4c), while the segment of residues 140-166 comprising of the two gear teeth is responsible for ring-ring interaction (Fig. 3d-f and Fig. 4d). As electron density missing often reflects the conformational heterogeneity arising from internal motions ^17^, these observation strongly suggest that the conformation of CsGSIb^Pen^ active site is highly dynamic, contrasting sharply to the conformationally largely homogeneous CsGSIb^Dec^. In support this, thermal shift assays show the melting temperature (T_m_) of wild-type CsGSIb is significantly lower than that of its mutants of I143L or EVK138DIQ, indicating of structural instability for the pentameric GSII (Fig. S10). We therefore conclude that the dramatically difference in the dynamic property of catalytic sites accounts for the distinct activities of pentameric and decameric CsGSIb. Taken together, our cryo-EM structures allow us to propose a dynamics-driven allosteric mechanism of how the GSII activity is regulated by changes in oligomeric state: (1) The active sites within isolated CsGSIb^Pen^ rings are highly disordered and the unstable catalytic environments render it catalytically inactive; (2) Upon stacking of two pentameric rings and formation of a decamer, the signals of interactions mediated by the gear teeth of each intervening subunit are allosterically propagated to the active sites, which reduce their conformational dynamics and in turn, unlock the catalytic potential of GSII (Fig. 4e).

**Fig. 4:**
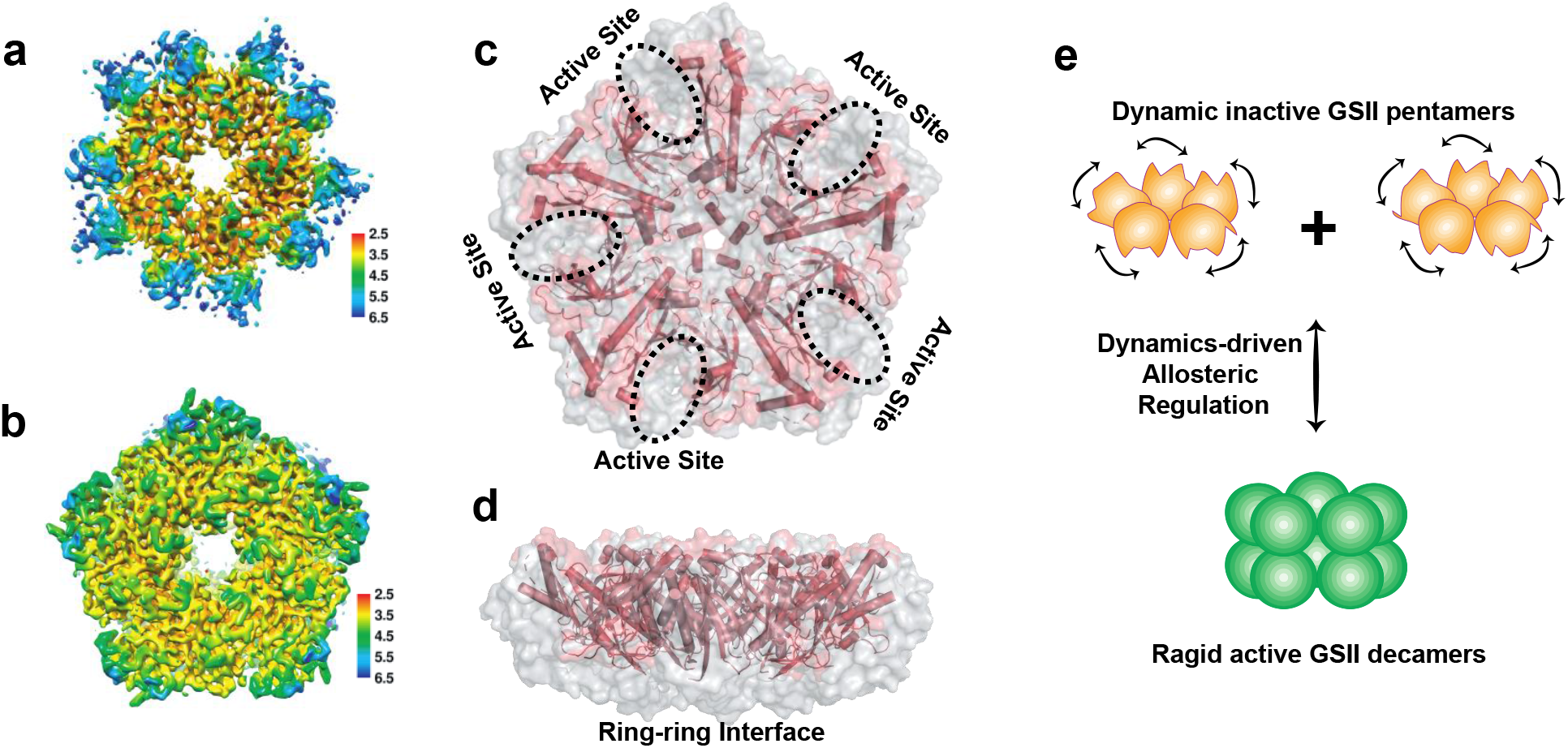
Cryo-EM structure of CsGSIb^Pen^ features high conformational flexibility. For simplicity, only one out three CsGSIb^Pen^ cryo-EM structures are presented here, and all three structures are shown in Fig. S9. **a**, Local resolution of the density map of CsGSIb^Pen^ indicates a decreased resolution near the edges of the pentamer ring. **b**, Local resolution of the density map of CsGSIb^Dec^ map. The conformational flexibility is apparent when the missing density at the rim of CsGSIb^Pen^, which yields a largely decagram-shaped map (**a**), is compared to the intact density of CsGSIb^Dec^ that displays a pentagon-shaped map (**b**). **c**-**d**, Superimposed of Cryo-EM structure of CsGSIb^Pen^ (pink) with that of CsGSIb^Dec^ (grey surface). **c**: Top-view; **d**: Side-view. The results reveal two major regions being highly disordered: the rim region including the catalytic center, and ring-ring interface. **e**, Proposed dynamics-driven allosteric model of CsGSIb.

### Activation of the GSII by the 14-3-3 scaffold protein

Having mechanistically established the allosteric coupling between GSII activity and its quaternary assembly status, we next asked whether there exist cellular factors that may regulate the CsGSIb activity, potentially via favoring its decamer assembly. 14-3-3 proteins are an important family of scaffold proteins that bind and regulate many key proteins involved in diverse intracellular processes in all eukaryotic organisms ^18–20^. In particular, self-dimerization of 14-3-3 proteins, which induces dimerization of their clients, plays a key role in its functional scaffolding and subsequent activity regulation ^18,19,21^. Moreover, it has been reported that 14-3-3 proteins act as an activator of GSs in various plants ^22–25^, although the detailed activation mechanism remains unclear. Based on these findings, here we tentatively provide the missing link in mechanistically assigning the role 14-3-3 proteins play in regulating GS activity: One protomer of the 14-3-3 protein recognizes one phosphorylated GSII pentamer, and its self-dimerization brings two pentamer rings in close proximity and therefore promotes decamer assembly, which, in turn, switches on the GS activity via allosteric rigidification of the catalytic sites (Fig. 5a). One prerequisite for this proposal is that, for the GS species whose activities being 14-3-3 protein-dependent, they must have an intrinsically weak decamer-forming propensity that is to be overcome by 14-3-3. In support of this, the GS from *Medicago truncatula*, whose activity is simulated upon binding to 14-3-3 protein ^24^, has been shown to exhibit a dynamic pentamer-decamer transition ^12^, similar to the CsGSIb presented here (Fig. 1d). Moreover, it has been shown that only the higher order complex of tobacco GS-2 that is bound to 14-3-3 is catalytically active ^25^.

**Fig. 5:**
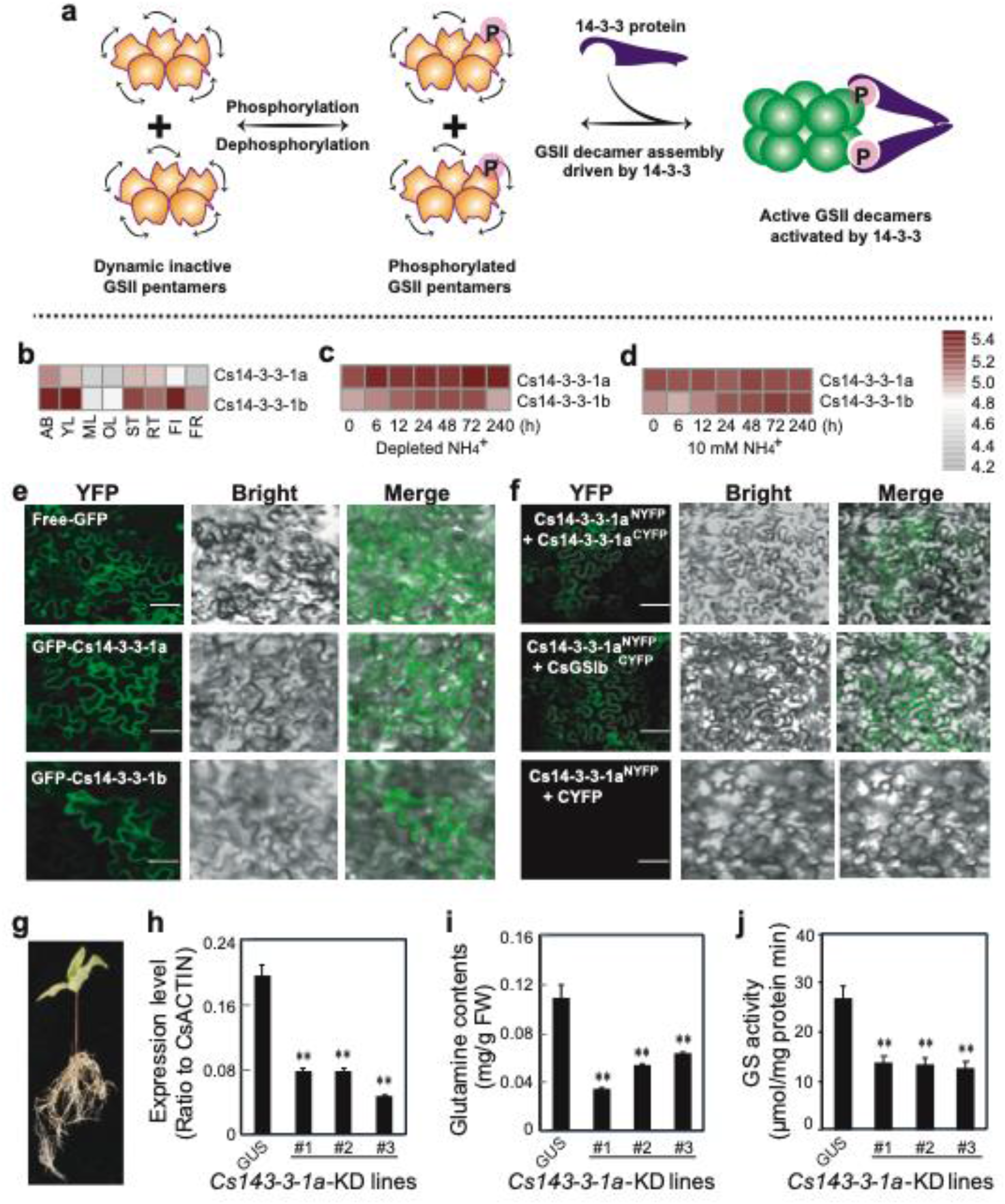
Activation of the CsGSIb by the 14-3-3 scaffold protein. **a**, Proposed working model of how 14-3-3 protein modulates the activity of GSIIs. P in pink sphere denotes the post-translational modification of phosphorylation. **b**, Expression patterns of *Cs14-3-3-1a* and *Cs14-3-3-1b* genes in various tea plant tissues. **c**, Induction of Cs14-3-3-1a and *Cs14-3-3-1b* genes by depletion of NH_4_^+^ from culture medium. **d**, Expression of *Cs14-3-3-1a* and *Cs14-3-3-1b* genes in tea plant roots fed with 10 mM NH_4_^+^. AB, apical buds of unopened leaves at the top of actively growing shoots; YL, first and second young leaves below the apical buds; ML, mature leaves geminated in the spring and harvested in the autumn; OL, old leaves at the bottom of tea tree plant; FL, flowers; FR, fruits of tea plants; ST, stem tissues at the 2nd and 3rd internodes; RT, roots. **e**. Subcellular localization of *Cs14-3-3-1a* and *Cs14-3-3-1a* fusion proteins in tobacco leaf epidermal cells. bar = 50 μM. **f**. BIFC assay of interaction among Cs14-3-3-1a, Cs14-3-3-1b and CsGSIIb in tobacco leaf cells. bar = 50 μM. **g**, Generation of tea plant transgenic hairy roots with RNAi knockdown(KD) of *Cs14-3-3-1a* gene. **h**, Expression of Cs14-3-3-1a in at least three tea plant transgenic hairy roots of *Cs14-3-3-1a-KD* compared with *GUS* roots. i, Glutamine contents in three tea plant transgenic hairy roots of *Cs14-3-3-1a-KD* compared with *GUS* roots. **j,** GS activity in three tea plant transgenic hairy roots of *Cs14-3-3-1a-KD* compared with *GUS* roots. All experiments were conducted at least three 3 independent experiments. At least 5 transgenic hairy roots for *Cs14-3-3-1a* and *GUS* genes were examined. Data are expressed as means ± s.d. Differences in two-tailed comparisons between transgenic lines and *GUS* controls were analyzed, **p < 0.01 in student’s t-test. See Methods for experimental details.

To further support the above proposal, we then explored whether the activity of the weak-decamer forming CsGSIb could also be regulated by 14-3-3 scaffold protein. Homology search against tea plant genome revealed several candidate tea plant Cs14-3-3 proteins (Fig. S11). Analysis of their coding genes’ expression patterns in tea plant tissues and in nitrogen assimilation or metabolism-related processes allowed us to identify *Cs14-3-3-1a* and *Cs14-3-3-1b* genes that displayed expression patterns highly similar to *CsGSI* genes (Fig. 5a and S12). Moreover, the expression levels of *Cs14-3-3-1a* and *Cs14-3-3-1b* genes were regulated upon changes in the availability of ammonia (Fig. 5b and 5c), the substrate of GS, suggesting both *Cs14-3-3-1a* and *Cs14-3-3-1b* are physiologically related to GS. We then examined the *in vivo* interactions between Cs14-3-3-1a and CsGSIb using the bimolecular fluorescence complementation (BiFC) technique, which is based on complementation between two non-fluorescent fragments of a fluorescent protein when they are brought together by interactions between proteins fused to each fragment ^26^. Cs14-3-3-1a or CsGSIb were fused in frame with N-terminal half of a yellow florescence protein (NYFP) or C-terminal half of a yellow florescence protein (CYFP), respectively, and expressed in tobacco leaf epidermal cells alone or in various combinations, such as CsGSIb ^CYFP^ alone or together with Cs14-3-3-1a^NYFP^. As expected, Cs14-3-3-1a and 1b could self-dimerize or form heterodimers in plant cells (Fig. 5e), consistent with 14-3-3 scaffold proteins adopting a dimeric structure ^18,19,21^. Importantly, formation of the fluorescent complex clearly demonstrated the interaction of Cs14-3-3-1a with CsGSIb (Fig. 5f). In order to further establish the functional relevance, we performed the RNA interference (RNAi) technique to knock down the transcript level of *Cs14-3-3-1a* gene in hairy roots of chimerical transgenic tea seedlings (Fig. 5h), and evaluated the impact on GS activity by measuring the contents of GS catalysis product, glutamine. We show that, along with the reduction in *Cs14-3-3-1a* transcript level, the glutamine contents (Fig. 5i), as well as the crude enzyme activity (Fig. 5j), were drastically reduced, indicating the 14-3-3 protein in *Camellia sinensis* functions as an activator molecule of CsGSIb. Further work is needed to elucidate the detailed mechanism of how *Cs14-3-3-1a* recognizes phosphorylated CsGSIb.

## Discussion

Proper assembly of individual protein units into functional complexes is fundamental to nearly all biological processes. Comparing to the oligomer assembled in relatively simple cyclic symmetry that contains only interfaces of subunits related by the rotational symmetry, protein complexes organized in dihedral symmetry, an extra step of assembly during evolution, possess interfaces that are related by both the rotational symmetry and the perpendicular two-fold axes. However, in many cases, the functional demand for such structural complexity remains poorly understood. Here, by using GS as a model system, we unveil a previously uncharacterized allosteric code buried in a supramolecular protein complex with dihedral symmetry, and show how dynamic packing of protein subcomplexes could build an extra allosteric control for activity modulation.

### Dynamics-driven allostery induced by assembly status transition

Allostery describes the mechanism that binding effector molecules at one site triggers a conformational or dynamic change at a distant site, thereby affecting protein activity. Allosteric regulation is a common mechanism to regulate protein function, playing critical roles in various cellular activities ranging from the control of metabolic mechanisms to signal-transduction pathways ^27^. Based on the data from Allosteric Database ^28^, to date more than 1,900 proteins have been defined as allosteric. Most allosteric modulators identified are small ligands or peptides, whereas in some rare cases the allosteric effects are induced by protein oligomerization in a rather simple system ^29^. Here we show that the oligomeric GS ring functions as positive modulator, the largest allosteric modulator identified so far to our knowledge; and the ring-ring association, which is motivated by 14-3-3 protein or other factors, leads to a transition of assembly symmetry from C5 to D5 and subsequently triggers allosteric activation.

We show here that the assembly geometry of GS plays a critical role in determining the protein functional motion properties (Fig. 4). Indeed, protein internal dynamics have been shown essential for functions; and allosteric proteins can be regulated predominantly by changes in their structural dynamics ^30–33^. The dynamics-driven allosteric mechanism presented here, in which the GSII activity is allosterically regulated by the change in conformational fluctuating properties of active sites via inter-ring communication, provides another fascinating example of the interplay between a protein’s dynamics and function.

### The allosteric mechanism of GSII offers a robust and tunable regulatory machinery

Being a key enzyme implicated in many aspects of the complex matrix of nitrogen metabolism, GS must be strictly regulated. Decades of efforts have been applied to understand how GS is controlled at gene, transcript and protein levels ^34^. Studies have demonstrated positive cooperativity of GS with regard to different substrates and cofactors, such as L-glutamate ^35^ and metal cations ^36^; and GS functions are regulated by multiple post-translational mechanisms including nitration, oxidative turnover and phosphorylation ^37^, and by the 14-3-3 protein ^22–25^. Our results reconcile with many of the above observations, and allow us to gain a more complete picture of how GS activity is regulated in cells by an exquisite machinery (Fig. 6). While the protein turnovers machineries to adjust cellular enzyme level can always provide means to modulate pentamer-decamer transitions and thus deactivation-activation conversion of GSII, we argue that phosphorylation-dephosphorylation processes, coupled with 14-3-3 binding and subsequent allosteric activation, may enable a more efficient regulatory way. When sufficient reaction products are available demanding low glutamine synthesis activity, GSII is kept in the dephosphorylated state by certain phosphatase and exists as isolated inactive single-ringed pentamers. In the physiological context of high demand of glutamine, phosphorylation of GSII by certain kinase prompts 14-3-3 protein binding, and the intrinsic dimerization property of 14-3-3 recruits two GSII pentamer rings in close proximity and in doing so, result in a rapid transition of quaternary assembly from the pentamer to decamer, and eventually enzymatic activation. In this manner, the poised GSII pentamer ring itself acts as a positive effector and the allosteric ring-ring association offers a great advantage of immediate response to precisely meet the ever-changing metabolic needs, whereas the reversible assembly-disassembly behavior enables a tunable mode for activity modulation. Indeed, the dynamic association-disassociation of GSII subcomplexes, a prerequisite for this modulatory machinery, have been widely observed in various species including humans ^38^, plants other than *Camellia sinensis* reported here ^12,15^ and fungi ^39^. Therefore, the dynamics-driven allostery shown here may represent a general regulatory machinery harnessed by many eukaryotes to ensure optimal utilization of nitrogen sources, and the infrastructure of fragile ring-ring contacts evolutionarily chosen by many eukaryotes offers a convenient and robust avenue for activity regulation.

**Fig. 6:**
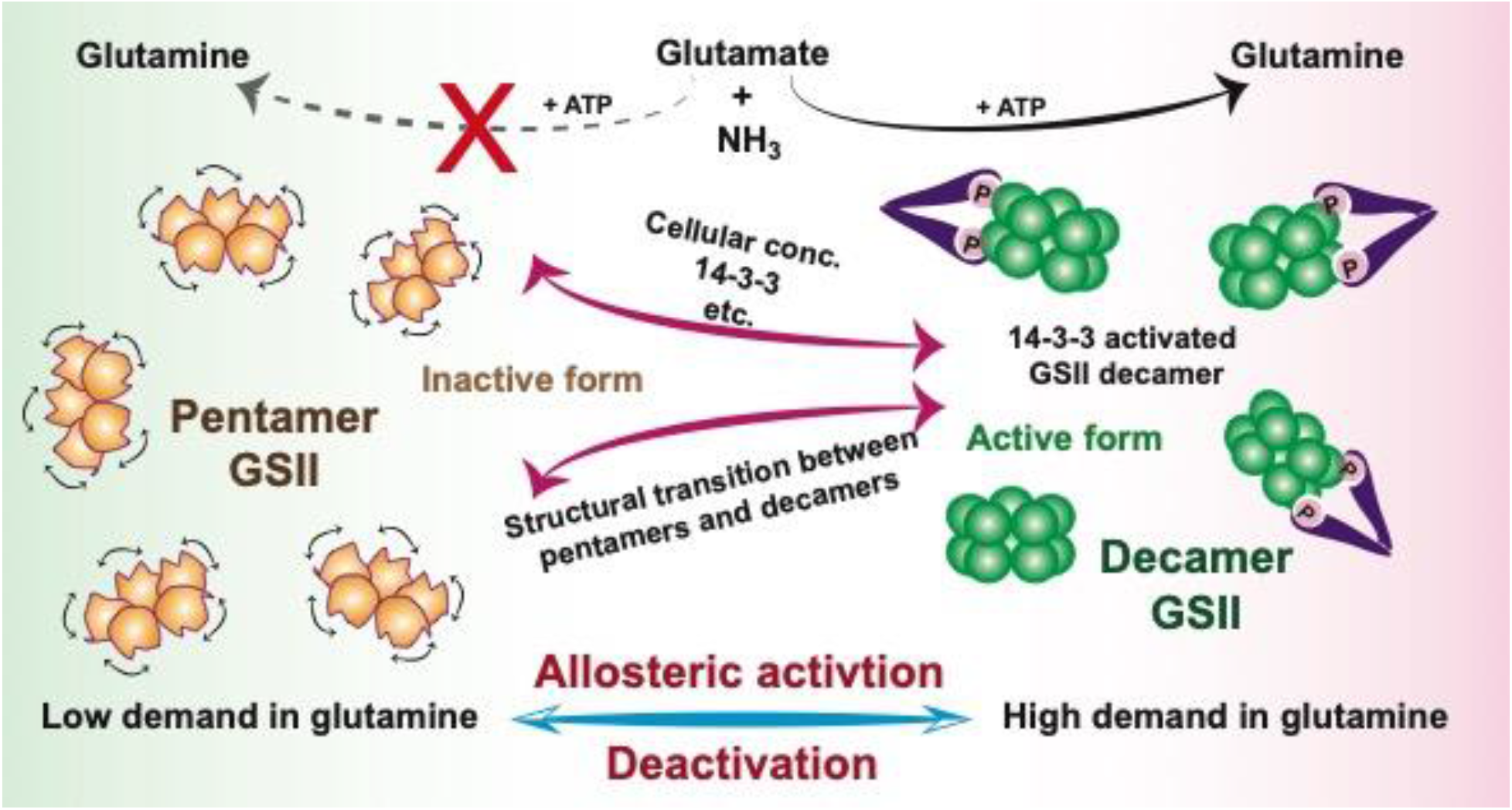
Schematic model of how activity of GSII is allosterically regulated in cells to meet metabolic needs. In low demand of glutamine, GSII is kept in the dephosphorylated state and exists as isolated inactive pentamers (orange); In the physiological context of high demand of glutamine, phosphorylation and subsequent binding to 14-3-3 (or by increase in cellular concentration) trigger a rapid transition of quaternary assembly of GSII from the pentamer to decamer (green), which, via the mechanism of dynamics-driven allostery, activates GSII.

### Practical implications

As a crucial enzyme to all living organisms, which is involved in all aspects of nitrogen metabolism, GS has emerged as an attractive target for drug design ^40^ and herbicidal compounds development, as well as a suitable intervention point for the improvement of crop yields ^41^. However, because the overall geometry of the active site is the most conserved structural element amongst GS enzymes ^4,7,11^, the traditional strategy of selective inhibition, which relies heavily on the subtle difference in the active sites from different species, has only achieved limited success. Thus, the regulatory mechanism discovered here will help guide the search for specific inhibitors of potential therapeutic interest. For example, inhibition of the GS in *Mycobacterium tuberculosis* has long been recognized as a novel antibiotic strategy to treat tuberculosis ^42–44^. Our result opens new possibilities to develop chemicals to target the drugable ring-ring interface region and specifically interrupt the interactions between two GS subcomplexes in pathogens or unwanted plants to develop new types of herbicide. Moreover, although overexpression of GS has been investigated extensively for decades with the goal of improving crop nitrogen use efficiency, the outcome has not been consistent ^41^. The modulatory “hot-spots” identified here, which mediate inter-ring communication and in turn stimulate GS activity, will guide engineering catalytically more powerful GSs for crops in which pentamer units only weakly associate, and thus increase plant nitrogen use efficiency and crop production.

## Methods

### Cloning, expression, and protein purification of GSs

The target genes encoding CsGSIb from *Camellia sinensis* (Genbank accession No. MK716208) and GmGSβ2 from *Glycine max* (Genbank accession No. NM001255403) were cloned into the pET-16b vector (Novagen) containing a His_6_-tagged-MBP tag followed by a tobacco etch virus (TEV) protease cleavage site at the N-terminus. All constructs were transformed into *E. coli* Rosetta (DE3) cells, which were cultured in Luria-Bertani (LB) medium at 37 °C supplemented with ampicillin (100 µg/ml) and chloramphenicol (35 µg/ml) to an OD600 ∼0.8. Cells were induced by the addition of isopropyl-β-D-1-thiogalactopyranoside (IPTG) to the concentration of 0.3 mM, and incubated for additional 16 hours at 18 °C. Cells were harvested by centrifugation at 5000g for 20 min and resuspended in lysis buffer (50 mM Tris-HCl, 500 mM NaCl, pH 8 and 1 mM PMSF).

Cells were subjected to a high-pressure homogenizer, named JN-Mini Pro Low-temperature Ultra-high-pressure cell disrupter (JNBIO) and then centrifuged at 50,000g for 30 min at 4℃. Proteins were initially purified using Ni Sepharose 6 Fast Flow resin (GE Healthcare). The protein tags were cleaved with His-tagged TEV-protease overnight at 4 °C while dialyzing against TEV cleavage buffer (50 mM Tris-HCl, 100 mM NaCl, 1 mM β-mercaptoethanol, pH 8). Cleaved sample was collected and run over Ni-NTA column to remove His-tagged TEV and protein tags. Flow-through was collected, concentrated and passed over Hiload 16/600 Superdex 200 column (GE Healthcare) in 50mM Tris-HCl, pH 7.4, 100 mM NaCl, 0.5 mM MgCl_2_ and 1.5 mM β-mercaptoethanol.

### Multi-angle light scattering (MALS) characterization

MALS was measured using a DAWN HELEOS-II system (Wyatt Technology Corporation) downstream of a GE liquid chromatography system connected to a Superdex 200 10/300 GL (GE Healthcare) gel filtration column. The running buffer for the protein samples contained 50 mM KPi (pH 7.0), 100 mM NaCl, 1 mM β-mercaptoethanol and 0.05 % NaN_3_. The flow rate was set to 0.5 mL min^−1^ with an injection volume of 200 μL, and the light scattering signal was collected at room temperature (∼23°C). The data were analyzed with ASTRA version 6.0.5 (Wyatt Technology Corporation).

### Glutamine synthetase activity assay

GS activity assay was performed described previously ^45,46^ and reactions were performed for 30 min at 37°C in 50mM Tris-HCl, pH 7.4, 100mM NaCl, 0.5 mM MgCl_2_ and 1.5 mM β-mercaptoethanol. Enzymatic activity comparison was conducted with 1 μM (monomer) enzyme in the presence of 0.5 mM NH_4_HCl, 2 mM L-glutamate, 0.5 mM ATP. Steady-state kinetic analysis was performed under the same conditions except with the variable concentration of ammonium chloride from 0.05 mM to 4 mM. Steady-state kinetic parameters were determined by double reciprocal Lineweaver-Burk plot for reactions that followed Michaelis-Menten kinetics. All experiments were repeated independently at least three times.

### Fluorescent dye-monitored thermal shift assays

Reactants containing 2uM CsGSIb (monomer) and 1000-fold diluted Sypro Orange in 50mM Tris-HCl, pH 7.4, 100mM NaCl and 1.5mM β-mercaptoethanol were performed using an iCycler thermocycler (Bio-Rad) as previous described ^13^. Briefly, CsGSIb in presence of various concentrations of the following ligands were tested, alone and in combination: 10mM glutamate, 20mM MgCl_2_ and 1mM ATP. The temperature of the reactions was increased from 20 to 90 °C in increments of 0.2 °C/12 s, coincident with a fluorescent measurement at each step. The wavelengths for excitation and emission were set to 490 and 575 nm, respectively. Fluorescence changes were monitored simultaneously with a charge-coupled device (CCD) camera. To obtain the temperature midpoint for the protein unfolding transition, Tm, a Boltzmann model was used to fit the fluorescence imaging data obtained by the CCD detector using the curve-fitting software GraphPad Prism 7.0.

### Analytical ultracentrifugation (AUC)

Sedimentation velocity experiments were carried out with a Proteomelab XL-A analytical ultracentrifuge (Beckman Coulter, USA) using a four-hole An-60 Ti analytical rotor. An aliquot of 410 μL of buffer (50 mM Tris-HCl, pH 7.4, 100 mM NaCl and 1.5 mM β-mercaptoethanol) as the reference and 400 μL of protein solution (0.1/0.25/0.5 mg⋅mL^−1^) were loaded into a double-sector cell. A centerpiece with a path length of 12 mm was used. The speed of rotor was 35,000 rpm. The operation temperature of rotor was 20 ℃. The time dependence of the absorbance at different radial positions was monitored at a wavelength of 280 nm by an UV–Vis absorbance detector, and the data were analyzed by the software SEDFIT (version 15.01b) using c(s) model to obtain the sedimentation coefficient distribution. Viscosity and density of the buffer solution were calculated by the Sednterp software.

### Single-particle cryo-electron microscopy data collection

Purified protein samples of CsGSIb (4 μL, 0.02 mg/mL) and GmGSβ2 (4 μL, 0.02 mg/mL) in 50 mM Tris-HCl, pH 7.4, 100 mM NaCl, and 1.5 mM β-mercaptoethanol were negatively stained with uranyl acetate 1% (w/v) on carbon-film 400 mesh copper grids. Samples were imaged using a FEI T12 operated at 120 keV with a 3.236 Å pixel size, 68,000× nominal magnification, and defocus range about 1.5 μm. For cryo-EM, 3 μL of CsGSIb (0.1 mg/mL and 0.5 mg/mL) and GmGSβ2 (0.1 mg/mL) were added onto glow-discharged Quantifoil R1.2/1.3 100 holey-carbon Cu grids with a Vitrobot Mark IV (Thermo Fisher Scientific). The grids were blotted for 3.5 s at 8 °C with 100% humidity, and then plunged frozen into liquid ethane cooled by liquid nitrogen. Cryo-grids were first screened on a FEI TF20 operated at 200 keV. Images of CsGSIb and GmGSβ2 were collected using Titan Krios G3i microscope (FEI) operated at 300 kV with a Gatan K2 Summit direct detection camera. Two datasets were acquired using the SerialEM in super-resolution mode with a nominal magnification of 29,000x, yielding a pixel sizes of 0.505 Å with a total dose of 51 e/ Å^2^. The defocus ranges were set from −1.6 μm to −2.3 μm.

### Cryo-electron microscopy image processing, 3D reconstruction, and analysis

All processing steps were performed using cryoSPARC ^47^. A total of 4,051 raw movie stacks acquired for CsGSIb and 1,777 raw movie stacks for GmGSβ2 were subjected to patch motion correction and patch CTF estimation. An initial set of about 500 particles were manually picked to generate 2D templates for auto-picking. The auto-picked particles were extracted by a box size of 512 pixel and then subjected to reference-free 2D classification. After particle screening using 2D and 3D classification, the final 355,289 particles for CsGSIb and 115,795 particles for GmGSβ2 were subjected to Ab-Initio Reconstitution and followed by 3D Refinement with C5 symmetry imposed. Four different conformational states were obtained for CsGSIb, resulting in a 3.3 Å density map for CsGSIb^Dec^, 3.5 Å density map for CsGSIb^Pen^ State I, 3.6 Å density map for CsGSIb^Pen^ State II, and 3.4 Å density map for CsGSIb^Pen^ State III. Only one major conformation was obtained with 2.9 Å density map for GmGSβ2. The global resolution of the map was estimated based on the gold-standard Fourier shell correlation (FSC) using the 0.143 criterion.

### Model building and structural refinement

Homology models of CsGSIb and GmGSβ2 were generated with the I-TASSER server ^48^ and docked into the cryoEM maps using UCSF Chimera ^49^. The sequences were mutated with corresponding residues in CsGSIb and GmGSβ2, followed by rebuilding in Coot ^50^. The missing residues of CsGSIb^Pen^ were not built due to the lack of corresponding densities. Real-space refinement of models with geometry and secondary structure restraints applied was performed using PHENIX ^51^. The final model was subjected to refinement and validation in PHENIX. The statistics of cryo-EM data collection, refinement and model validation are summarized in Table S1.

### Different nitrogen treatments for hydroponically grown tea cuttage seedlings

Two-year-old hydroponic tea cuttage seedlings were grown in a greenhouse at 20-25°C until new tender roots emerged. These healthy tea seedlings were then transferred into hydroponic solutions with different nitrogen sources, namely, 0 mM NH_4_^+^ (Shigeki Konishi solution), 5 mM NH_4_^+^ (Shigeki Konishi solution with 5 mM ammonium nitrogen), 10 mM NH_4_^+^ (Shigeki Konishi solution with 10 mM ammonium nitrogen), and a control (Shigeki Konishi solution alone). All of these tea seedling root samples were cleaned and collected in liquid nitrogen after treatment for RNA analysis.

### RNA isolation and qRT-PCR analysis

Tea plant tissues or root materials were ground in liquid nitrogen into fine powders for total RNA extraction with an RNA extraction kit (Tiangen Biotech Co., Ltd.) according to the manufacturer’s instructions. RNA quality and purity were assessed by a NanoDrop 2000 spectrophotometer (Thermo Scientific). The integrity of the RNA samples was rapidly checked by 1.0 % agarose gel electrophoresis. The total RNA was reverse-transcribed to single-stranded cDNAs using SuperScript III reverse transcriptase (Invitrogen) according to the manufacturer’s instructions. qRT-PCR analysis was performed using cDNA synthesized by the Prime Script RT Reagent Kit (TaKaRa). Each qRT-PCR was conducted in a 20-μL reaction mixture containing 2 μL of diluted template cDNA, 0.4 μL of each specific primer, 10 μL of SYBR Premix Ex-Taq (TaKaRa), and 7.2 μL of H_2_O. All qRT-PCR assays were performed on the Bio-Rad CFX96 fluorescence-based quantitative PCR platform. The program used was as follows: 95°C for 5 minutes; 40 cycles of 95°C for 5 s for denaturation and 60°C for 30 s for annealing and extension; and 61 cycles of 65°C for 10 s for melting curve analysis. All experiments were independently repeated three times, and relative expression levels were measured using the 2−ΔCt method.

### Subcellular localization of CsGSIs and Cs14-3-3-1a&1b

Construction of the Cs14-3-3-1a-GFP, CsGSI-1b-GFP, and Cs14-3-3-1b-GFP fusions were performed using gateway recombination systems. The corresponding ORFs for CsGSIb and Cs14-3-3-1a, 1b were subcloned into pK7WGF2 in frame with a GFP tagged at the N-terminus. Determination of the subcellular localization of these GFP fusions was performed using tobacco leaf infiltration as previously described (Zhao et al., 2011). Briefly, the pK7WGF2-Cs14-3-3-1a-GFP, pK7WGF2-Cs14-3-3-1b-GFP, and pK7WGF2-CsGSIb-GFP plasmids were transformed into *A. tumefaciens* strain EHA105, and selected positive colonies harboring these constructs were used for plant transformation by infiltration. *Acetosyringone*-activated Agrobacterium cells were infiltrated into the *Nicotiana benthamiana* leaves leaf abaxial epidermal surface, and the tobacco plants were grown at room temperature for 3 days before imaging. Imaging of these GFP fusion proteins was performed using a confocal microscope with a 100× water immersion objective and appropriate software. The excitation wavelength was 488 nm, and emissions were collected at 500 nm.

### In planta interaction between CsGSIb with Cs14-3-3 proteins with Bimolecular Fluorescent Complimentary (BIFC)

The ORFs of Cs14-3-3-1a, Cs14-3-3-1b, and CsGSIb were amplified and subcloned into pCAMBIA1300-eYFPN (YFP N-terminal portion) and pCAMBIA1300-eYFPC (YFP C-terminal portion) (CAMBIA) by the in-fusion technology. The resulting constructs were transformed into A. tumefaciens strains GV3101, which were infiltrated into Nicotiana benthamiana leaves individually or in different pair combinations. A Leica DMi8 M laser scanning confocal microscopy system was used for fluorescence observation, according to the method described previously ^52^. If the fluorescence signal could be detected with any interaction pair, the pair of half YFP-fusion proteins should interact.

### Knockdown of Cs14-3-3-1a in tea plant hairy roots

Approximately 400 bp of the gene-specific fragments from Cs14-3-3-1a were amplified and subcloned into the final RNA interference (RNAi) destination vector pB7GWIWG by BP and LR clonase-based recombination reactions (Invitrogen). The resulting binary vectors pB7GWIWG-Cs14-3-3-1a were transformed into *A. rhizogenes* strain ATCC 15834 by electroporation. The selected positive transformants harboring pB7GWIWG-Cs14-3-3-1a were used to transform 3-month-old tea seedlings, which were pretreated with acetosyringone. The positive transgenic hairy root lines were verified with qRT-PCR for examination of transgene expression. At least three independent hairy root lines were selected for further analysis.

### Determination of free amino acids in tea plant samples

The free amino acids in tea plant samples were analyzed by using an amino acid analyzer (L-8900, Hitachi) according to manufacture instruction. The free amino acids were extracted from 120 mg leaves with 1 mL of 4% sulfosalicylic acid in water bath sonication for 30 min and then centrifuged at 13,680 x g for 30 min. The debris was re-extracted once again and the supernatants from two extractions were combined as previously described ^53^. The supernatants were filtered through a 0.22 µm Millipore filter before analysis. A mobile phase containing lithium citrate for amino acid derivatization and UV–Vis detection at 570 and 440 nm were used in the Hitachi High-Speed Amino Acid Analyzer system. The flow rates were set at 0.35 mL/min for the mobile phase and 0.3 mL/min for the derivatization reagent. The temperature for separation column was set to 38 °C, and for the post-column reaction equipment was maintained at 130 °C. The temperature of the autosampler was kept at 4 °C. The peak areas of amino acids were quantified in comparison with the amino acid standards.

### GS activity assay from plant samples

To determine the total GS activity, 100 mg of frozen plant samples were grounded into fine powder in liquid nitrogen. Samples were homogenized in extraction buffer (50 mmol/L Tris-HCl, pH 8.0, 2 mmol/L MgSO_4_, 4 mmol/L dithiothreitol, and 0.4 mmol/L sucrose). Plant extracts were centrifuged at 13,680 x g (4 °C) for 25 min and the supernatants of extracts were analyzed for the soluble protein content using the Bradford assay. GS activity was determined after incubating the enzyme extracts in a reaction buffer (100 mmol/L Tris-HCl, 80 mmol/L MgSO_4_, 20 mmol/L sodium glutamate, 80 mM NH_4_OH, 20 mmol/L cysteine, 2 mmol/L EGTA and 40 mmol/L ATP) at 37 °C for 30 min ^54^. A stop solution containing 0.2 mol/L Trichloride acetic acid, 0.37 mol/L FeCl_3_ and 0.6 mol/L HCl was added; and the absorbance of enzyme reactions at 540 nm was recorded. A standard curve was made in an identical way for calculation of the specific enzyme activity.

## Supplementary Data & Figures

### *Camellia sinensis* CsGSIb exhibits a pentamer-decamer dynamic equilibrium in solution

To further characterize the low propensity of CsGSIb for decamer forming, we carried out SEC-MALS measurements at various concentrations. As shown in Fig. S1a, increase in the concentration of CsGSIb sample injected favored association of pentamers towards formation of decamer, and vice versa, suggesting a dynamic nature of the interaction between two pentameric rings. Moreover, a series of SEC-MALS measurements of CsGSIb at different salt concentrations were performed to decipher the driving force mediating the inter-ring contact. While addition of salt of NaCl in the regime of low salt concentrations (from 0 to 300 mM) provoked the dissociation of decamer into pentamers first, further increase in salt concentration (from 300 to 500 mM) tended to facilitate re-association of pentameric subcomplexes to some extent (Fig. S1b), suggesting a mixture of electrostatic and hydrophobic interactions is responsible for attaching of two pentameric rings.

To probe the effects of various ligands on the stability of CsGSIb, a series of fluorescent dye-monitored thermal shift assays were carried out as described previously^13^. As shown in Fig. S1c, addition of magnesium ions or its combination with the nonhydrolyzable ATP analog AMPPNP resulted in increases in the melting temperature (T_m_) of CsGSIb with ∼6 or ∼10 °C, respectively, while the presence of the substrate of glutamate showed no apparent effect on T_m_, consistent with previous observations ^13^. In contrast, as evidenced by SEC-MALS measurements (Fig. S1d), the presence of the above ligands showed no appreciable effect on the decamer-forming properties. These observations collectively indicate that while binding to substrate or cofactor rigidifies the structural organization within individual pentamer rings, the inter-ring assembly of GSII is, at least in vitro, not substrate-induced.

**Fig. S1:**
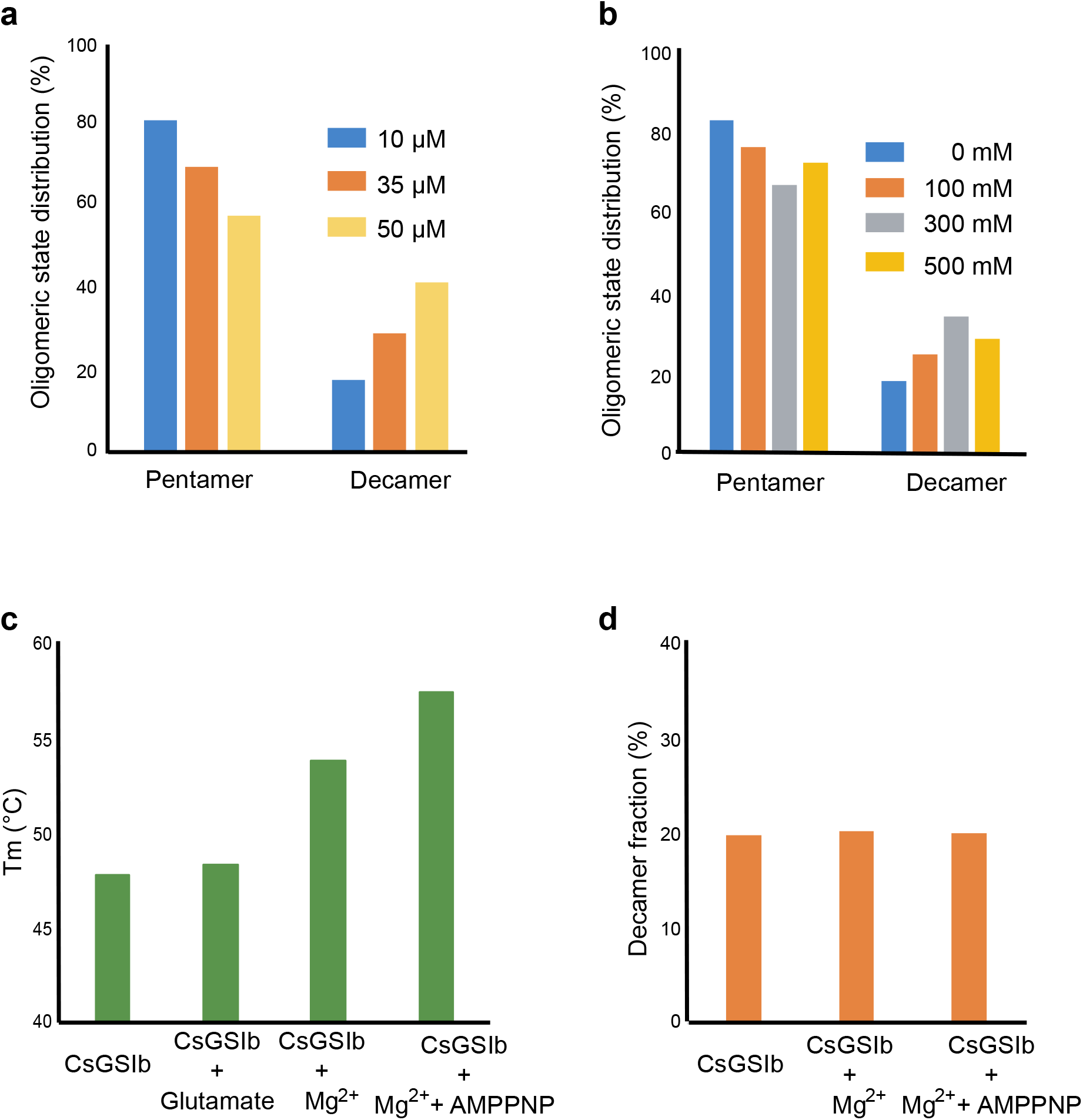
CsGsIb exhibits a pentamer-decamer dynamic equilibrium in solution. Using SEC-MALS to reveal the dependence of CsGsIb assembly status on: **a**, protein concentrations; **b**, salt concentrations. **c**, Melting temperatures of CsGsIb in presence of various substrates as measured by differential scanning fluorimetry. **d**, Effects of various substrates on quaternary assembly status of CsGsIb as measured by SEC-MALS.

**Fig. S2:**
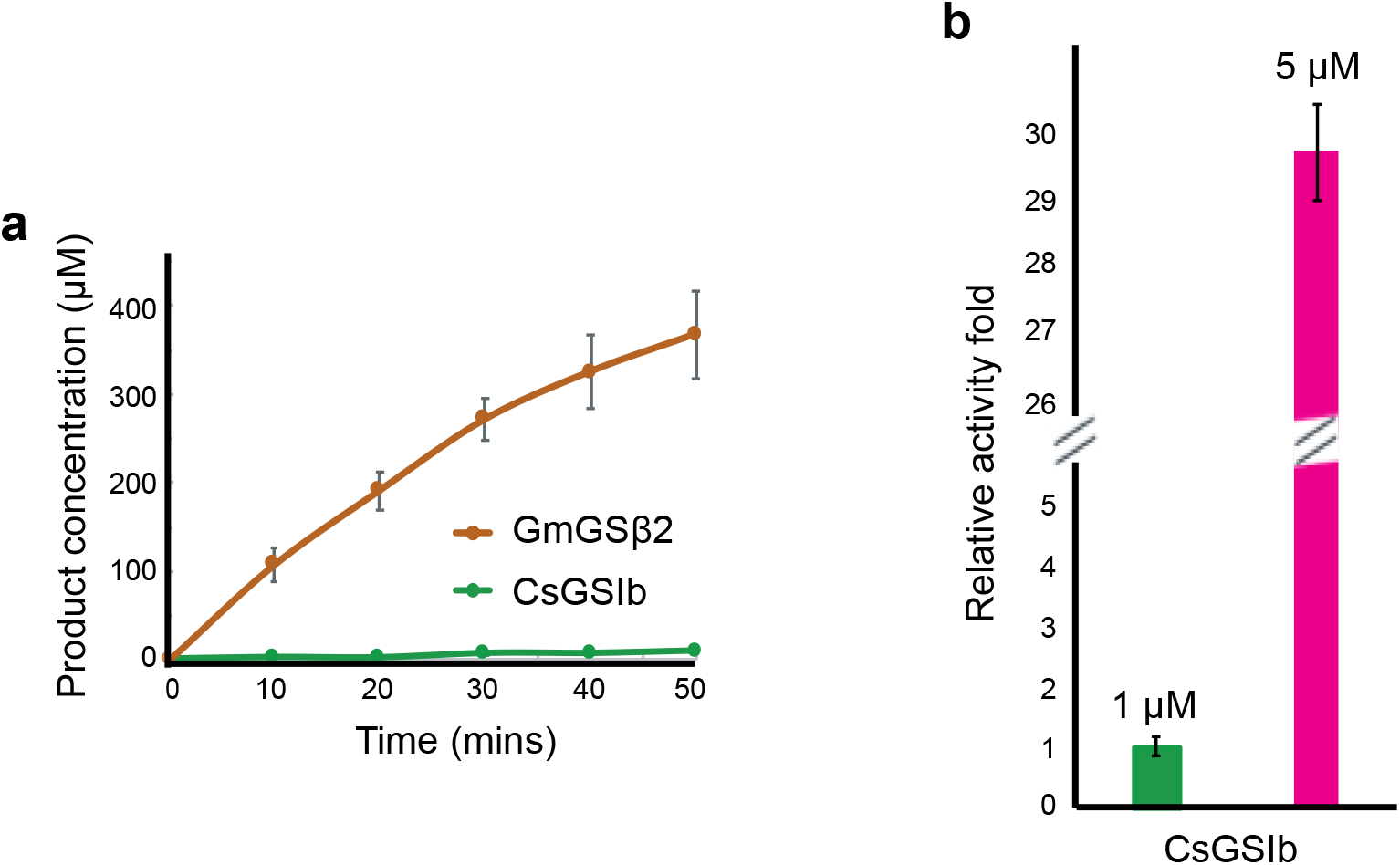
Enzymatic characterizatioin of GmGSβ2 and CsGSIb. **a**, Time courses of GS activity for GmGSβ2 (orange) and CsGSIb (green). **b**, GS activity of CsGSIb demonstrates significant concentration-dependence. Green bar: 1 μM enzyme; Pink barL 5 μM enzyme, Enzyme concentrations are shown as monomer concentration. Reaction conditions are same as that in Fig. 2. All experiments were repeated three times and data are shown as means ± s.d.

**Fig. S3:**
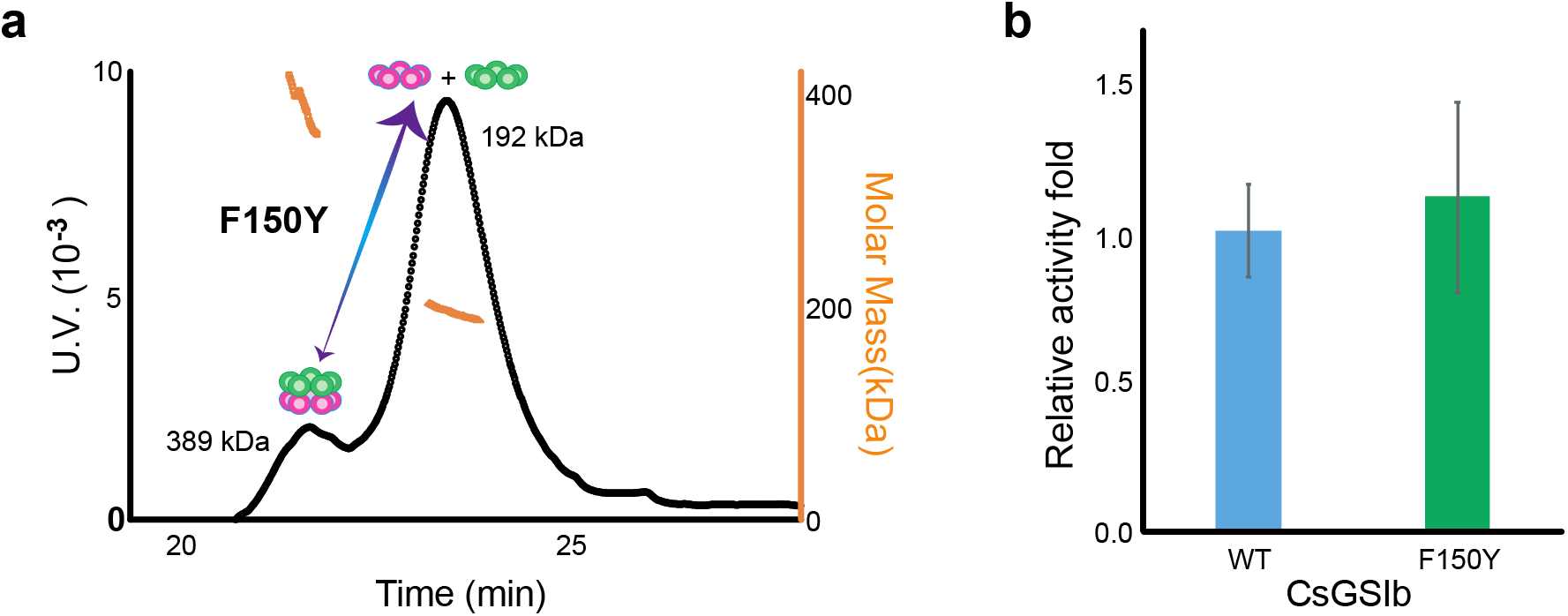
Mutation effects of F150Y on the quaternary assembly property and enzyme activity of CsGsIb. **a**, SEC-MALS profile of CsGsIb F150Y mutant reveals no change in the quaternary assembly property upon mutation. **b**, Wild type CsGsIb and F150Y demonstrate similar GS activities. Activity assays were performed three times in the condition same as that in Fig.2. Data are shown as means ± s.d.

**Fig. S4:**
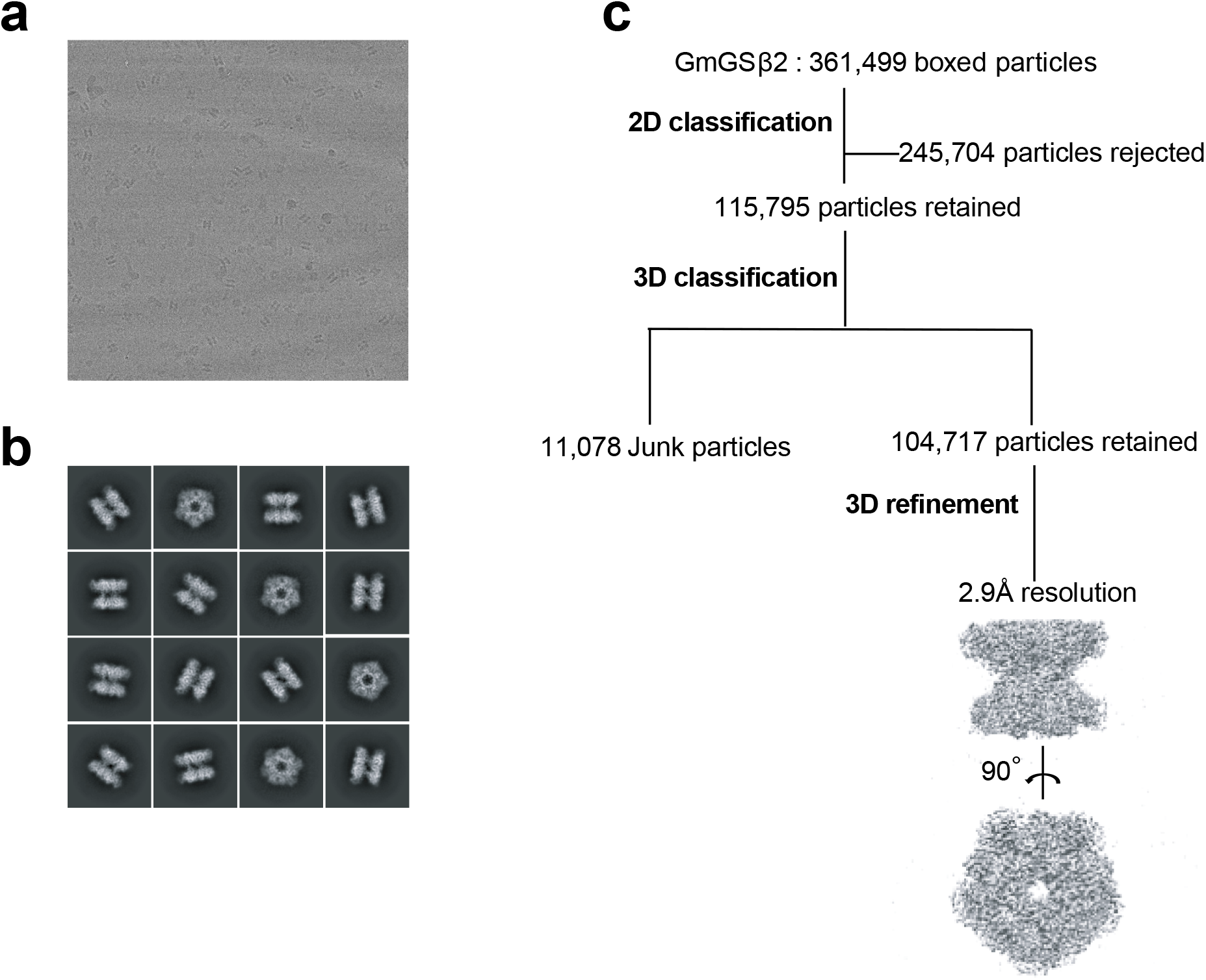
Cryo-EM anaysis of GmGSβ2. **a,** Representative cryo-EM micrograph; **b,** Subset of representative, reference-free 2D class averages; **c**, Data processing workflow.

**Fig. S5:**
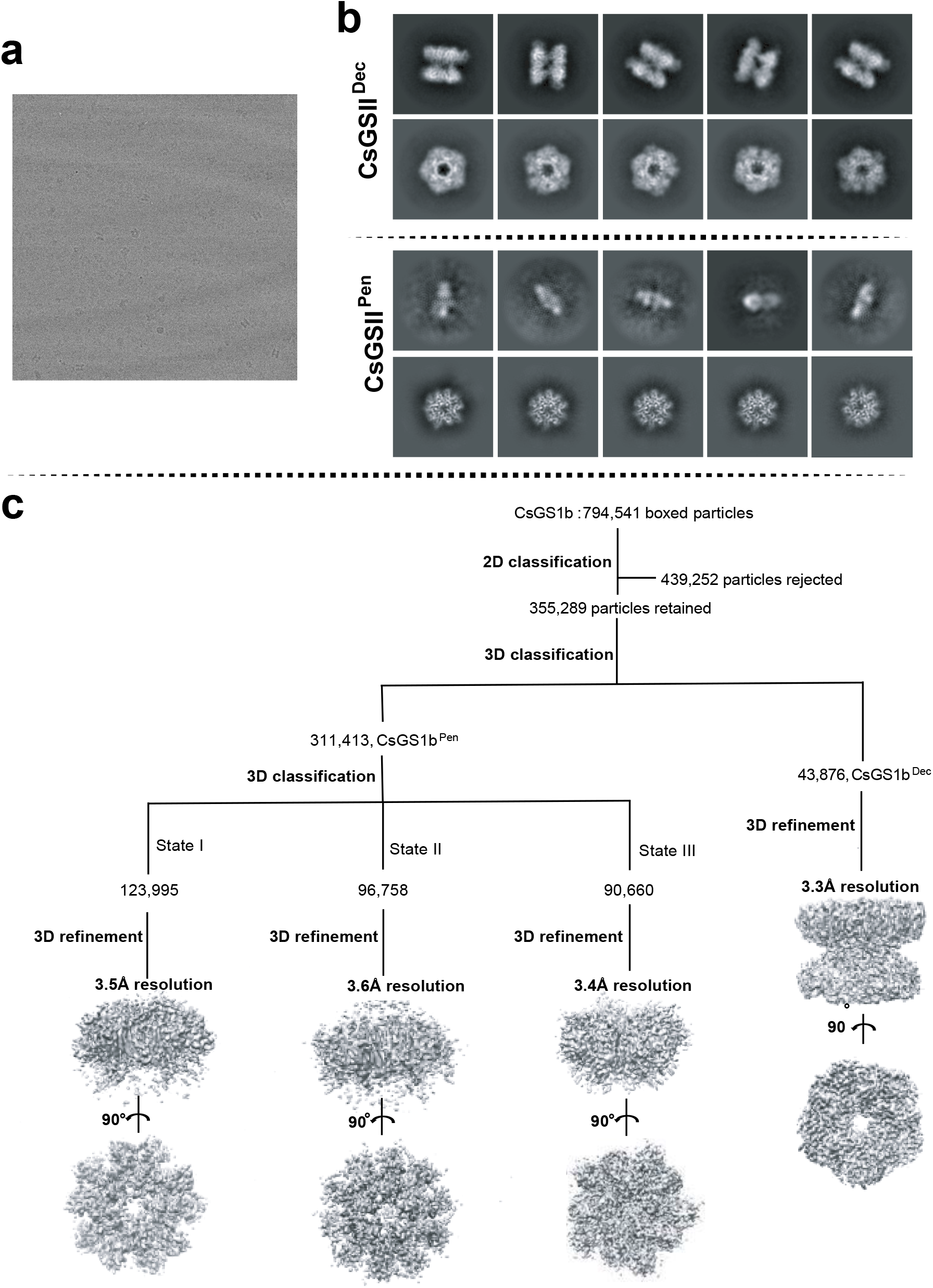
Cryo-EM anaysis of CsGSIb. **a,** Representative cryo-EM micrograph; **b,** Subset of representative, reference-free 2D class averages; **c**, Data processing workflow.

**Fig. S6:**
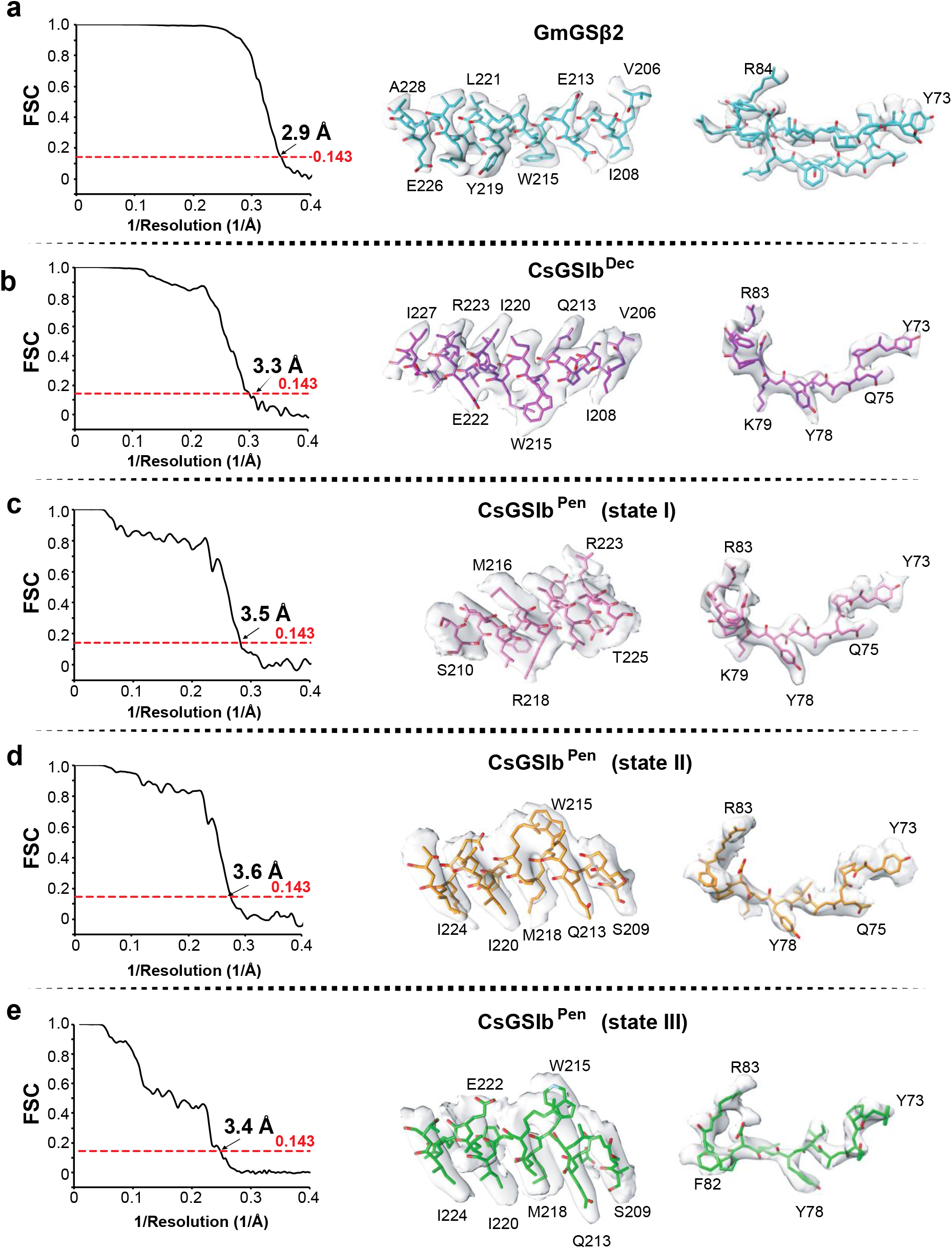
Validation of cryo-EM structures. **a-e:** GmGSβ2, CsGSIb^Dec^, and three states of CsGSIb^Pen^, respectively. Left: Gold-standard FSC plots generated from cryoSPARC; Right: View of model fitted in representative density.

**Fig. S7:**
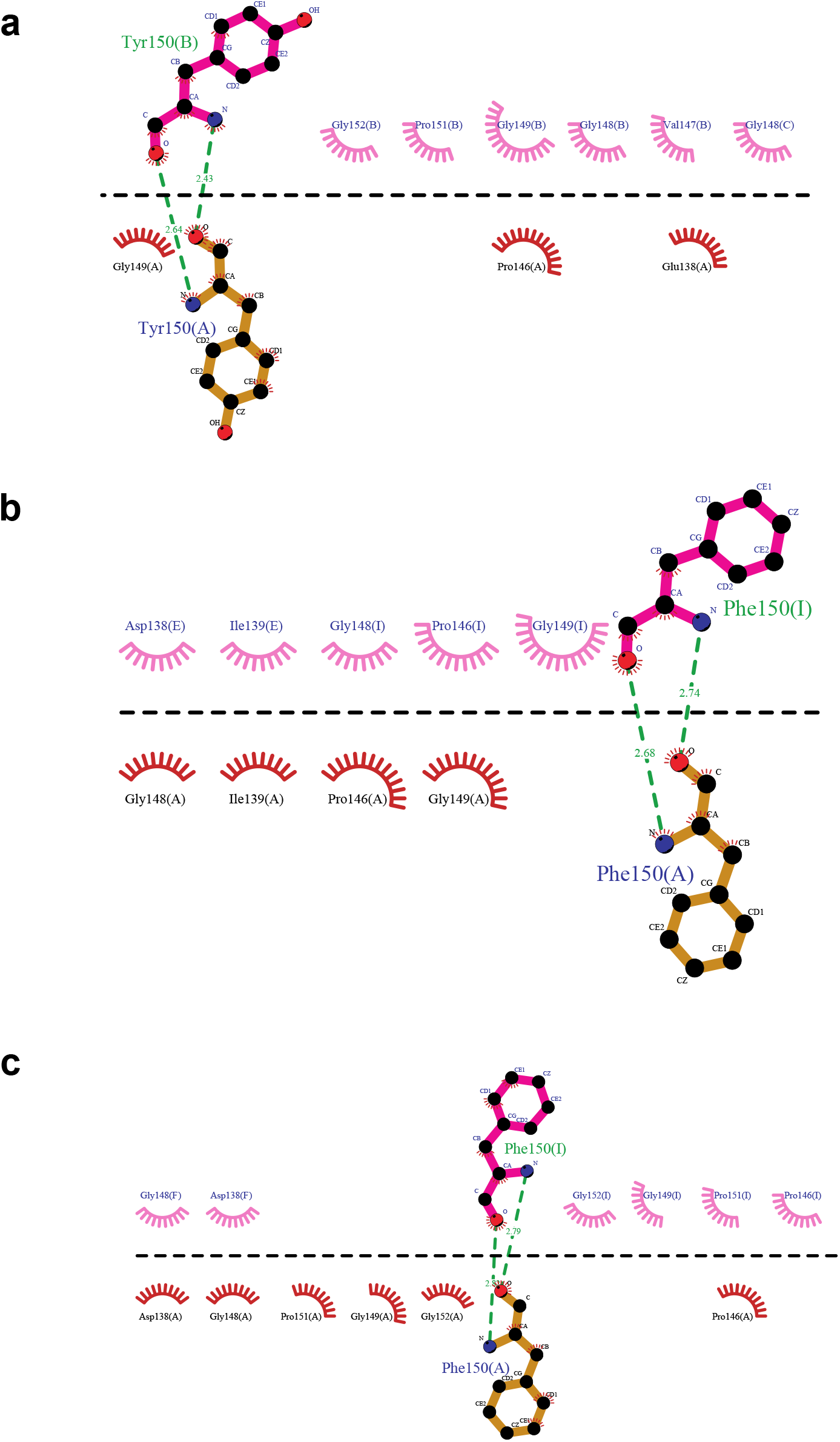
Detailed analysis of interactions bewtween two GSII pentamric rings using the program of Ligplot. **a**, CsGSIb; **b**, GmGSβ2; and **c**, Maize GS using the pdb code of 2d3a.

**Fig. S8:**
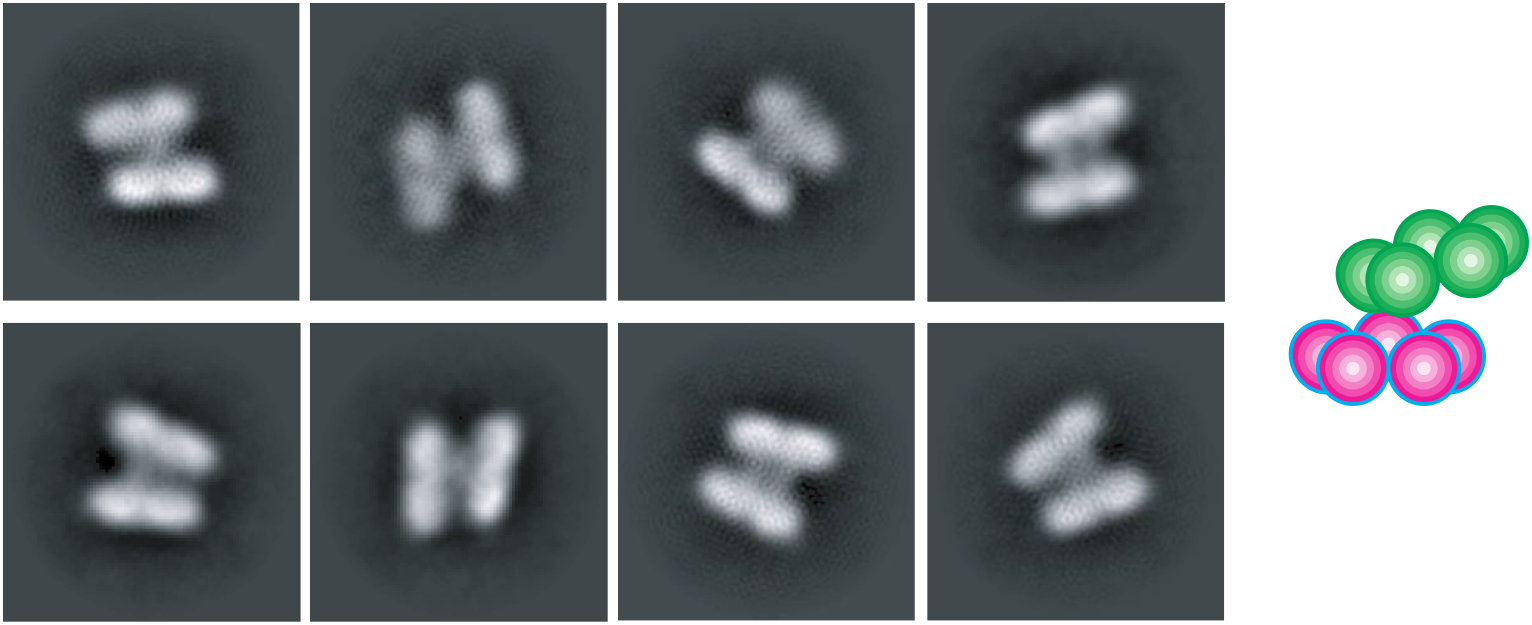
Swinging motion of CsGsIb rings. Two-dimensional class averages of CsGsIb particles reveals a few class of particles in which two pentameric rings are no longer parallel. This swinging motion of the rings with respect to each other is likely to be owing to flexibility of the inter-ring connections. Right: A schematic representation of the averages is shown for clarity.

**Fig. S9:**
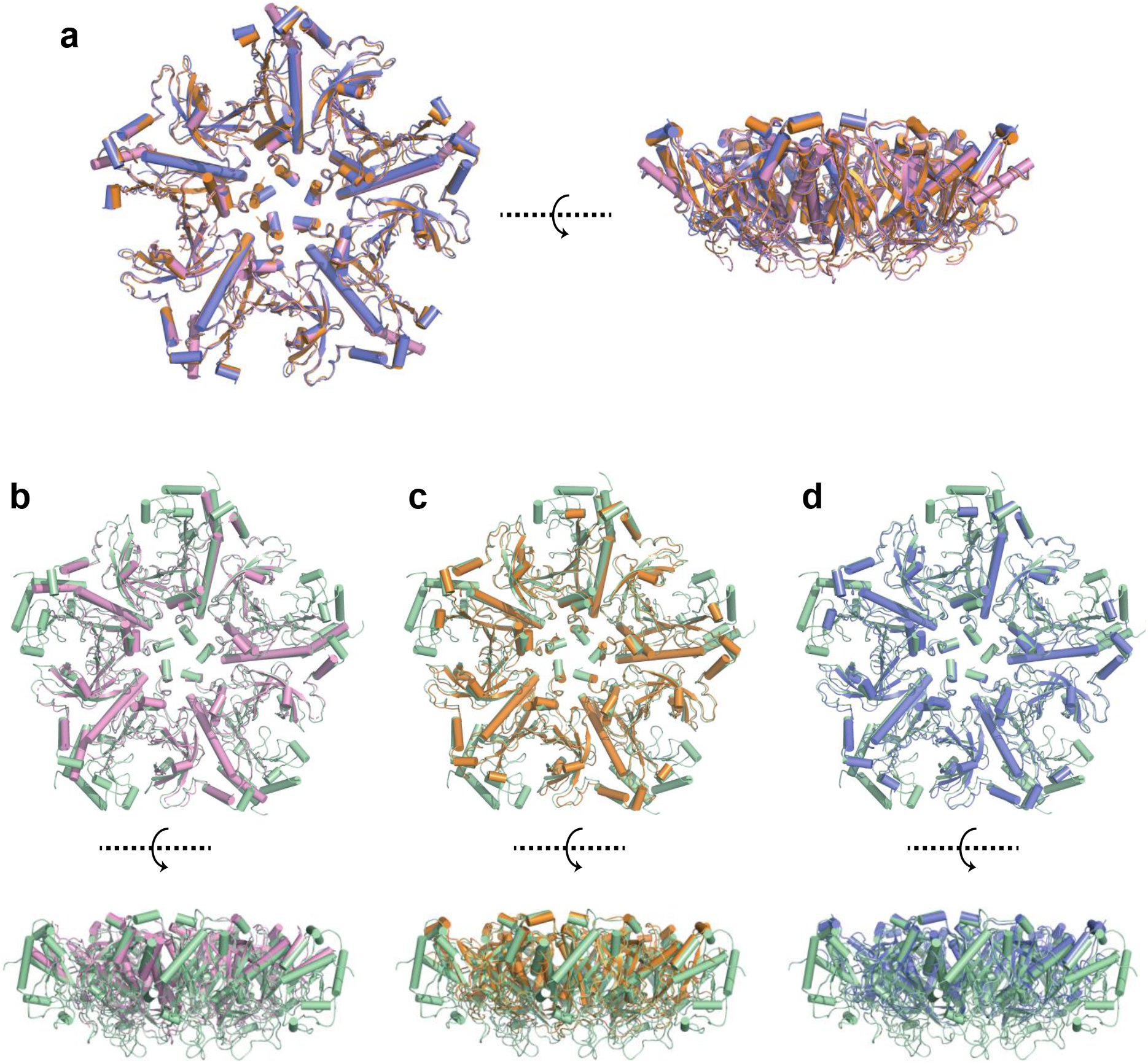
Three cryo-EM structures of CsGSIb pentamers in isolation. **a,** Structure allignment of three CsGSIb^Pen^ structures determined using Cryo-EM. Left: Topview; Right: Sideview. These three stuctures, colored in golden, pink and pruple, respectively, are highly similar to each other, with only a few structural variations at the peripheral regions. **b-d**, Structure comparison of three structures of CsGSIb pentamer in isolation (colored the same as in **a**) with that in the context of decamer (in color of green). Upper: Topview; Lower: Sideview. Note a large portion is missing in the structure of CsGSIb^Pen^ arising from electron density missing, indicating those regions are highly dynamic.

**Fig. S10.**
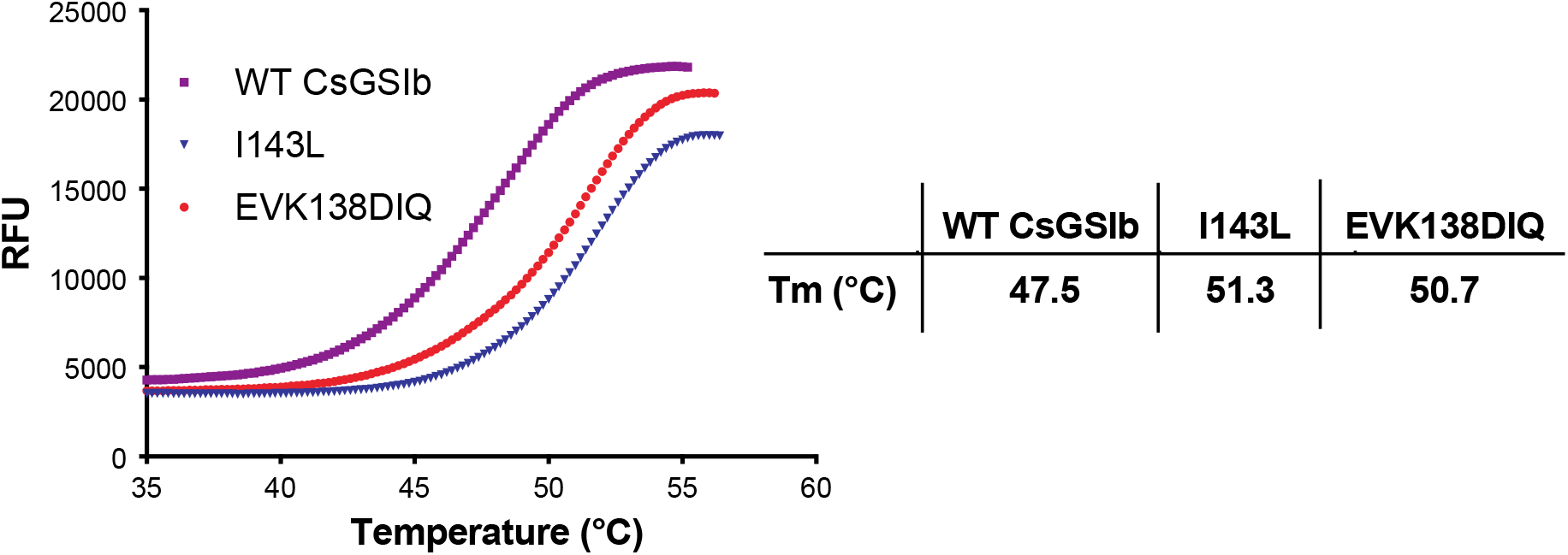
Thermal shift assays for CsGSIb and mutants. The thermal stabilities of wild type CsGsII and mutants were analysed by measuring SYPRO Orange dye fluorescence over a temperature ranging from 35 to ∼55 °C using a real-time PCR thermocycler. Left: Representative unfolding curves; Right: Derived meting temperatures. RFU: Relative fluorescence unit. The lower value of meting temperature for WT CsGSIb indicates structural instablity for the wild type CsGSIb.

**Fig. S11.**
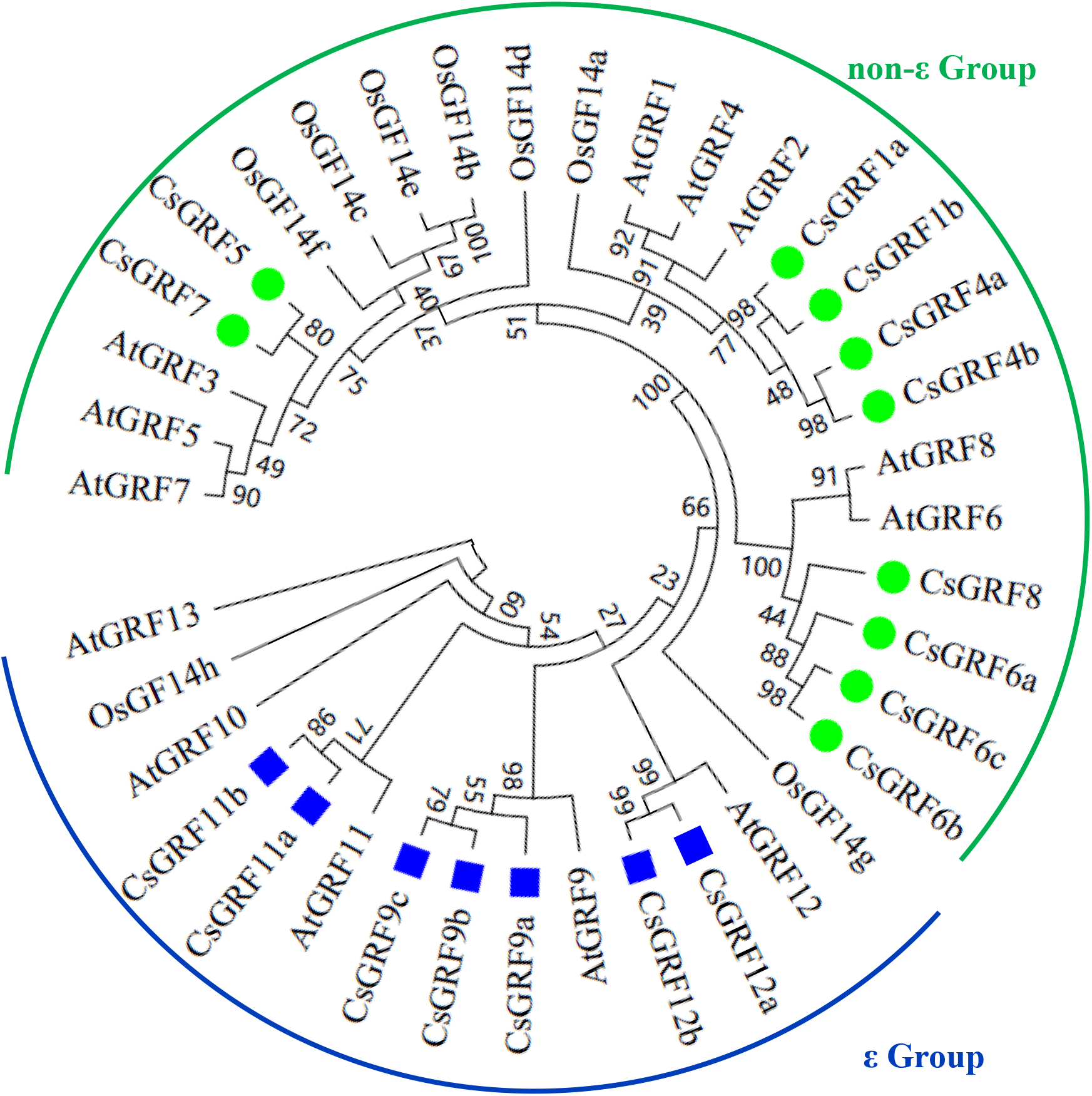
Phylogenetic analysis of Cs14-3-3s from *Camellia sinensis* genome as compared with 14-3-3s from *Arabidopsis* and rice. Amino acid sequences were aligned by Clustal W. MEGA 6.0 software was used to construct the phylogenetic tree by the NJ method with 1000 bootstrap replicates. They are divided into two groups of non-ε Group and ε Group.

**Fig. S12.**
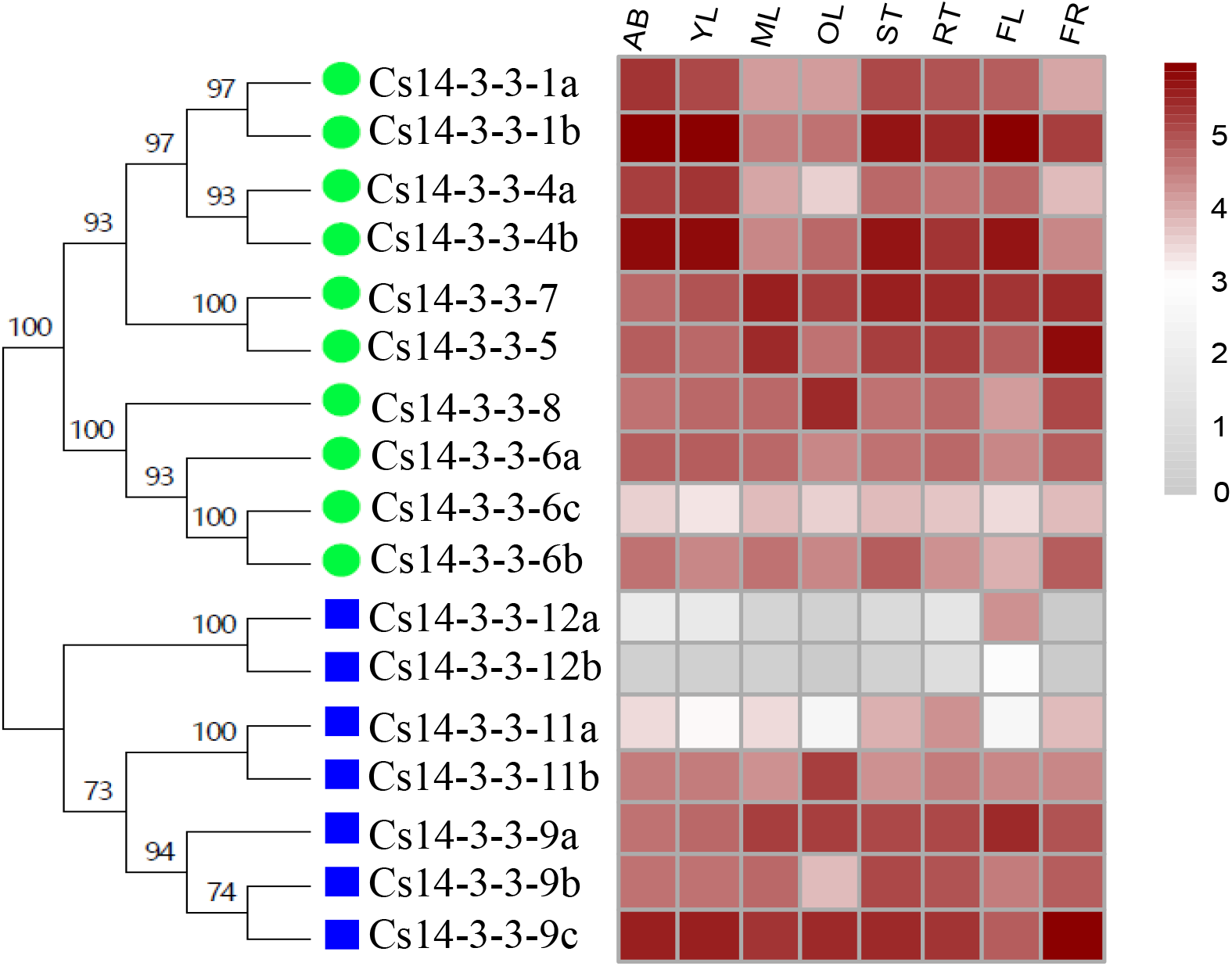
Expression of *Cs14-3-3* genes in eight tissues of *Camellia sinensis* plants., including AB, apical bud refers to unopened leaves on the top of activity growing shoots; YL, young leaf includes the first and second leaf below the apical bud; ML, mature leaf is for these geminated in the spring and are harvested in the autumn; OL, old leaf for these in the bottom of tea tree plant; FL, Flower; FR, fruit of tea plants, ST, Stem for the 2nd, 3rd internodes; RT, roots, were retrived from RNA Sequencing data. Expression levels were calculated using Log10 (FPKM).

**Table S1.** Cryo-EM data collection, refinement, and validation statistics.

|  | CsGS1b<sup>Dec</sup><br>(PDB 7V4I) | CsGS1b<sup>Pen</sup> (I)<br>(PDB 7V4J) | CsGS1b<sup>Pen</sup> (II)<br>(PDB 7V4K) | CsGS1b<sup>Pen</sup> (III)<br>(PDB 7V4L) | GmGSβ2<br>(PDB 7V4H) |
| --- | --- | --- | --- | --- | --- |
| **Data collection** |  |  |  |  |  |
| **Magnification** | 29000 x | 29000 x | 29000 x | 29000 x | 29000 x |
| **Voltage (kV)** | 300 | 300 | 300 | 300 | 300 |
| **Electron exposure**<br>**(e<sup>-</sup>/Å<sup>2</sup>)** | 51 | 51 | 51 | 51 | 51 |
| **Defocus range (μm)** | -1.6 ~ -2.3 | -1.6 ~ -2.3 | -1.6 ~ -2.3 | -1.6 ~ -2.3 | -1.6 ~ -2.3 |
| **Pixel size (Å)** | 0.505 | 0.505 | 0.505 | 0.505 | 0.505 |
| **Reconstruction and Model composition** |  |  |  |  |  |
| **Symmetry imposed** | D5 | C5 | C5 | C5 | D5 |
| **Chains** | 10 | 5 | 5 | 5 | 10 |
| **Nonhydrogen atoms** | 27680 | 9850 | 9260 | 8966 | 27340 |
| **Protein residues** | 3560 | 1275 | 1195 | 1140 | 3520 |
| **Refinement** |  |  |  |  |  |
| R.m.s. deviations |  |  |  |  |  |
| **Bond lengths (Å)** | 0.003 | 0.003 | 0.004 | 0.004 | 0.004 |
| **Bond angles (°)** | 0.585 | 0.670 | 0.660 | 0.696 | 0.589 |
| **Model-to-map fit (CC)** | 0.74 | 0.52 | 0.64 | 0.57 | 0.82 |
| **Map resolution (Å)** | 3.3 | 3.5 | 3.6 | 3.4 | 2.9 |
| **FSC threshold** | 0.143 | 0.143 | 0.143 | 0.143 | 0.143 |
| **Validation** |  |  |  |  |  |
| **MolProbity score** | 1.91 | 2.28 | 1.98 | 2.24 | 1.63 |
| **Clashscore** | 10.27 | 19.49 | 17.27 | 17.81 | 7.07 |
| **Rotamers outliers (%)** | 0.00 | 0.00 | 0.20 | 0.00 | 0.00 |
| **Ramachandran plot** |  |  |  |  |  |
| **Favored (%)** | 94.46 | 91.84 | 96.33 | 91.87 | 96.37 |
| **Allowed (%)** | 5.54 | 8.16 | 3.67 | 8.13 | 3.63 |
| **Outliers (%)** | 0.00 | 0.00 | 0.00 | 0.00 | 0.00 |

## Notes

### Competing Interest Statement

The authors have declared no competing interest.

